# Integrated high-confidence and high-throughput approaches for quantifying synapse engulfment by oligodendrocyte precursor cells

**DOI:** 10.1101/2023.08.24.554663

**Authors:** Jessica A. Kahng, Andre M. Xavier, Austin Ferro, Yohan S.S. Auguste, Lucas Cheadle

## Abstract

Oligodendrocyte precursor cells (OPCs) sculpt neural circuits through the phagocytic engulfment of synapses during development and in adulthood. However, precise techniques for analyzing synapse engulfment by OPCs are limited. Here, we describe a two-pronged cell biological approach for quantifying synapse engulfment by OPCs which merges low-and high-throughput methodologies. In the first method, an adeno-associated virus encoding a pH-sensitive, fluorescently-tagged synaptic marker is expressed in neurons *in vivo.* This construct allows for the differential labeling of presynaptic inputs that are contained outside of and within acidic phagolysosomal compartments. When followed by immunostaining for markers of OPCs and synapses in lightly fixed tissue, this approach enables the quantification of synapses engulfed by around 30-50 OPCs within a given experiment. In the second method, OPCs isolated from dissociated brain tissue are fixed, incubated with fluorescent antibodies against presynaptic proteins, and then analyzed by flow cytometry. This approach enables the quantification of presynaptic material within tens of thousands of OPCs in less than one week. These methods extend beyond the current imaging-based engulfment assays designed to quantify synaptic phagocytosis by brain-resident immune cells, microglia. Through the integration of these methods, the engulfment of synapses by OPCs can be rigorously quantified at both the individual and populational levels. With minor modifications, these approaches can be adapted to study synaptic phagocytosis by numerous glial cell types in the brain.

## Introduction

The construction of neural circuits during brain development occurs in a stepwise fashion, beginning with the establishment of an overabundance of nascent synaptic connections *in utero*. This early phase of synapse formation is followed by the strengthening and maintenance of a subset of synapses, coinciding with the large-scale elimination of synapses that are transient or dispensable for mature brain function^1,2^. The removal of synapses during development is essential for ensuring that the proper number and organization of synaptic connections persist across the lifespan. Although synapse elimination has predominantly been studied during development, this process occurs in the mature brain as well, likely to facilitate the remodeling of established circuits in response to extrinsic stimuli^3–5^. Thus, synapse elimination is a fundamental mechanism driving the development and plasticity of the brain.

Over the past ten years, the elimination of excess synapses has been shown to involve the phagocytic engulfment and degradation of presynaptic inputs by glia, non-neuronal cells of the brain. While prior studies have mainly focused on the engulfment of synapses by microglia^6–8^, we recently discovered a new role of oligodendrocyte precursor cells (OPCs) in engulfing synapses both during postnatal development and in the adult mouse brain^9^. Along with recent complementary studies^10,11^, this finding shed light on the ability of OPCs to sculpt brain circuits and influence brain function beyond the production of mature oligodendrocytes. Thus, OPCs are multi-functional brain cells with key roles in circuit connectivity and function across the lifespan. These discoveries have revealed a need for precise techniques tto investigate how OPCs perform their non-canonical functions, including the engulfment of synapses.

### Development of the Approach

The discovery that OPCs engulf synapses in the brain was facilitated by the development of methods for visualizing and quantifying synaptic material within OPCs. An important requirement for these experiments was the ability to identify synaptic structures within OPCs with a high degree of confidence. Unlike other glia, OPCs receive direct synaptic inputs from neurons. Therefore, differentiating between synaptic contacts at the surface of OPCs from synaptic material that is internalized is particularly important and may lead to unique challenges compared to other phagocytic glia^12,13^. For instance, one obstacle to accurately identifying internalized synapses within OPCs is the limitation of spatial resolution inherent to confocal microscopy. Although the enhanced resolution afforded by super-resolution microcopy techniques can help identify engulfed synapses, these methods are typically neither as readily available nor user-friendly as confocal microscopes. Furthermore, even with improved resolution, these approaches tend to be low-throughput, which may miss phenomena only quantifiable when observing a larger population of cells as a whole.

In the current protocol, we describe two complementary approaches: one that utilizes standard confocal imaging and quantifies OPC engulfment of synaptic material in a high-confidence but low-throughput manner, and a second approach that uses flow cytometry to analyze synaptic engulfment by OPCs in a high-throughput manner (Figure 1). The first method takes advantage of the fundamental observation that, when a cell phagocytoses extracellular material, that material is shuttled into phagolysosomes (PLs) where it is degraded. These organelles are highly acidic, allowing us to engage a presynaptically anchored, pH-sensitive construct to identify whether OPC-ingested synaptic material resides within a PL. These adeno-associated viral (AAV) sensors include <u>p</u>robe for <u>syn</u>aptic <u>dig</u>estion (pSynDig)^9^ and ExPre^4^, both of which fuse the presynaptic protein Synaptophysin to an mCherry (red) and an eGFP (green) fluorophore. When synapses are intact and at physiological pH, pSynDig+ synapses fluoresce in both red and green channels. However, when inputs reside within acidic compartments, they lose green signal at a higher rate than the red signal due to eGFP’s limited pH resilience^9^. Thus, eGFP-negative inputs in OPCs are most likely inputs that have been phagocytosed and are in the process of being degraded.

**Figure 1.**
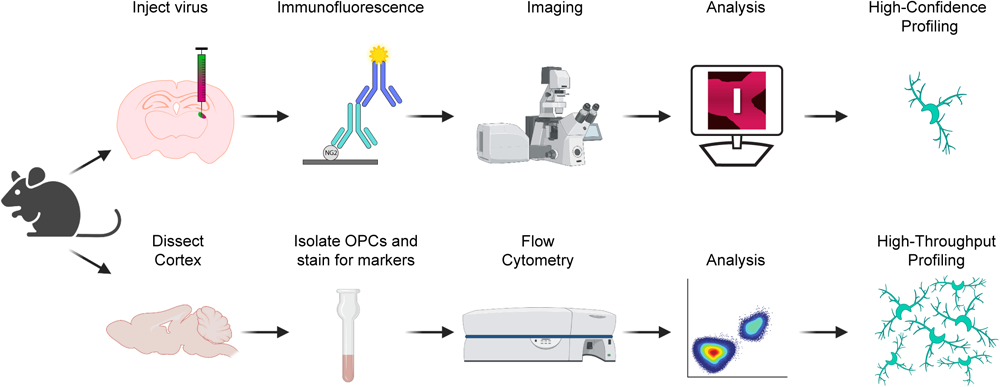
Overview of the experimental design. The two-pronged approach to quantify synapse engulfment by oligodendrocyte precursor cells (OPCs) in the mouse brain involves an imaging-based strategy to analyze a relatively limited number of OPCs at a high level of confidence (top) and a flow cytometry-based approach to profile synapses within OPCs at a populational level (bottom).

In approach 1, pSynDig is delivered to a brain region such that neurons in that region project fluorescently labeled presynaptic inputs that terminate in the region on interest. In this protocol we focus on the infection of neurons in the dorsal lateral geniculate nucleus (dLGN) of the thalamus and the subsequent labeling of their inputs in primary visual cortex. However, this approach can be adapted to a broad range of circuits. To localize pSynDig+ axonal inputs within OPCs in visual cortex, about three weeks after thalamic viral infection, the brain tissue is lightly perfusion-fixed, harvested, and subjected to immunofluorescence staining with antibodies against (1) the OPC marker NG2 and (2) markers for presynaptic inputs such as VGLUT2. The co-stained tissue is then imaged, with three-dimensional Z-stacks acquired using either a standard or Airyscan confocal microscope at Nyquist settings. Imaging at Nyquist settings ensures that the sampling frequency of the image is at least twice the highest frequency present in the specimen. This results in a more accurate representation of the specimen which improves the accuracy of the quantitative measurements. After image acquisition, OPCs and synaptic inputs are reconstructed in the software package Imaris (Figure 2a). A filtering-based algorithm is applied to identify, with high confidence, synaptic inputs that are localized within a given OPC. These inputs can be analyzed either by quantifying the volume of synaptic signal within an OPC normalized to the OPC’s volume (yielding an *engulfment score*), or by measuring the eGFP-to-mCherry signal ratio within the cell (Figure 2b). These strategies allow for the rigorous quantification of the amount of engulfed synaptic material within about 30 - 50 OPCs in each experiment.

**Figure 2.**
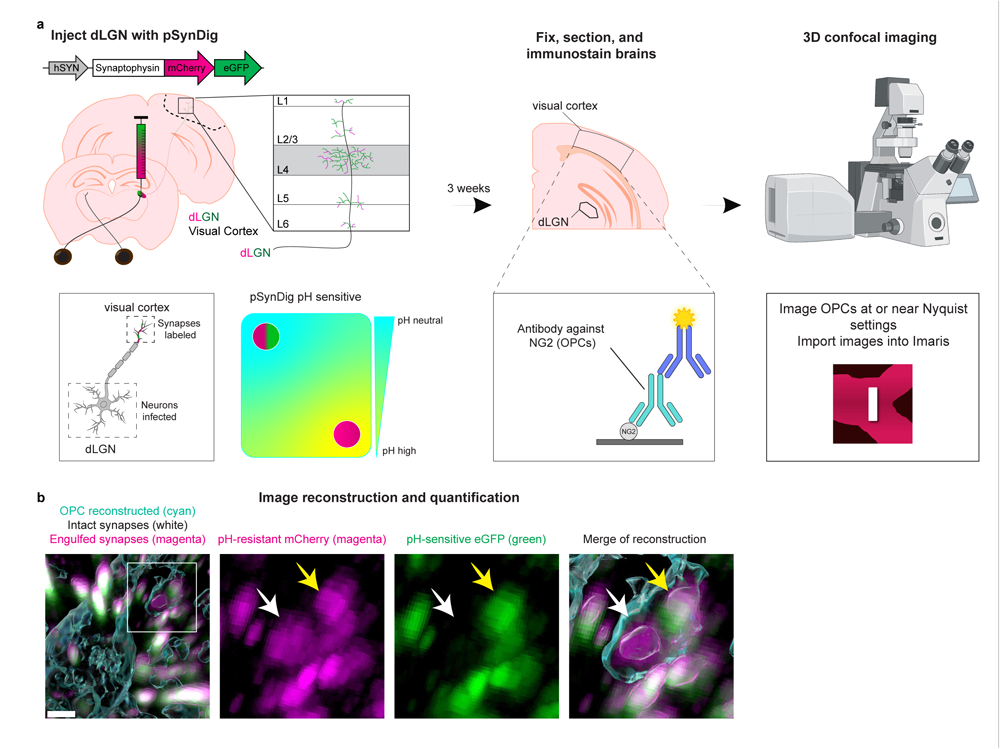
Low-throughput imaging-based engulfment assay. (a) Schematic illustrating the imaging-based approach for quantifying synapses within OPCs. First, pSynDig is injected into the dorsal lateral geniculate nucleus (dLGN) of the thalamus of a mouse to label presynaptic inputs in visual cortex. After three weeks, the brain is harvested, sectioned, and immuno-stained for markers of OPCs (e.g., NG2). Markers of synapses (e.g., VGLUT2) can also be stained for at this point. Finally, volumetric 3D images of immuno-stained visual cortex are taken on a confocal microscope and imported into Imaris. (b) In Imaris, OPCs (cyan) and pSynDig (intact synapses containing eGFP and mCherry, white; digested synapses not containing eGFP, magenta) are reconstructed. Yellow arrow, intact synapse. White arrow, digesting synapse. The filtering-based approach is then applied to quantify the volume of synaptic material within OPCs at a high level of confidence. Scale bar, 4 µm.

Approach 1 describes an imaging-based method that more accurately quantifies OPC engulfment of synapses than the current strategies used for other glia types. However, this method is low-throughput, and transcriptomic and functional studies have indicated that OPCs may have multiple cell states^14–17^. Thus, low-throughput methods are likely insufficient to fully understand the nature of OPCs engulfment of synaptic material, including whether these cells engulf in a homogenous or heterogeneous manner. Therefore, we have established a second approach to quantify the relative expression of presynaptic markers within individual OPCs across a pool of tens of thousands of OPCs using flow cytometry (Figure 3). In this approach, OPCs are first dissociated from micro-dissected cortical tissue and then are isolated by density through centrifugation together with other glial cells, particularly microglia. These cells then go through two rounds of staining, one that stains for extracellular markers specific for OPCs and another that stains for presynaptic proteins located intracellularly (Figure 3a). Post-staining, these cells are then analyzed by flow cytometry. This allows for the quantification of the presynaptic protein content within OPCs at the populational level (Figure 3b). Although this method does not allow for the direct visualization of the intracellular content, it is a fast and reliable complement to imaging-based analyses with the benefit of quantifying synaptic material within OPCs in a high-throughput manner^18^. Because the imaging-based and flow cytometry-based approaches are highly complementary, their combination allowed us to define a heterogeneous function for OPCs in engulfing synapses in the developing and mature brain^9^.

**Figure 3.**
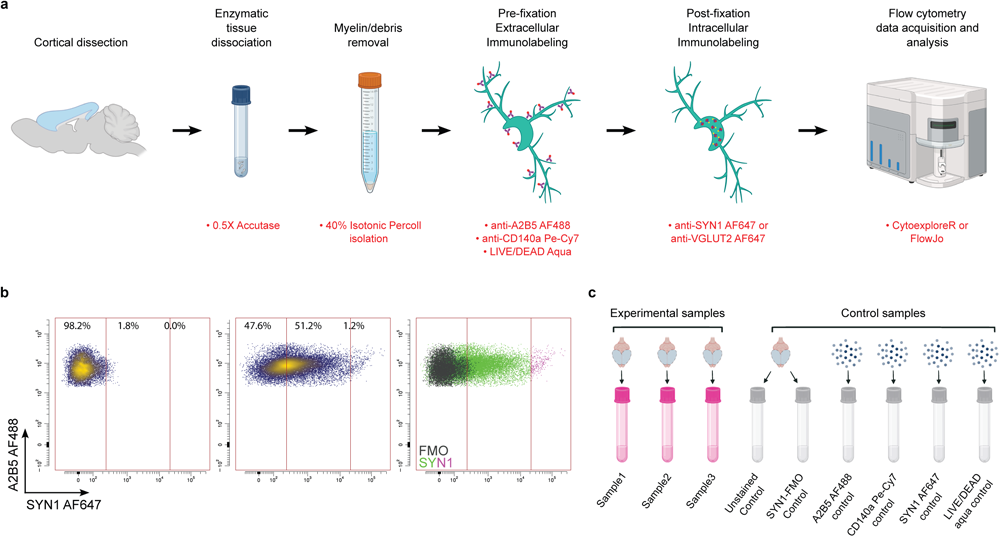
High-throughput flow cytometric assay. (a) Schematic depicting the experimental workflow for analyzing presynaptic protein content in OPCs. Following euthanasia, brain tissue is collected, and cortical dissection is performed. The tissues are then incubated with enzyme and subjected to manual tissue dissociation and homogenization. The resulting homogenate is centrifuged in isotonic percoll to remove myelin and debris, and OPCs are stained for extra-and intracellular protein targets. Flow cytometry data is acquired and analyzed using cytoexploreR or FlowJo software. (b) Flow cytometry plot showing the presence of the presynaptic marker SYN1 within OPCs. The fluorescence minus one (FMO) control sample is included for reliable gating strategy. Data points are colored based on the median fluorescence intensity of SYN1 marker within the OPCs (gray, FMO control; green, OPCs incubated with SYN1; magenta, OPCs containing a particularly large amount of synaptic material). (c) Experimental design for a single flow cytometry run.

### Comparison with other methods

Historically, synapse engulfment by glial cells has been quantified using an imaging-based approach in fixed tissue simply termed the ‘engulfment assay’^6^. In this assay, synaptic inputs are labeled with antibodies against presynaptic proteins (e.g., VGLUT2) while cell volumes are labeled with antibodies against microglia (e.g. IBA1, P2ry12). Three-dimensional Z-stacks are then acquired on a confocal microscope, after which the labeled elements are reconstructed using a specialized software program called Imaris. Imaris uses a masking algorithm to quantify the volume of a cell in question that is occupied by synaptic material. This value, which is normalized to cell volume, serves as a read-out for the engulfment activity of a given cell.

While this approach was innovative and remains widely used, there are three major limitations of this method. (1) The spatial resolution typically achieved on a standard confocal microscope is often not fine enough to confidently discriminate whether a synaptic input is within, in contact with, or just very close to a cell. (2) These experiments are relatively low-throughput, making it difficult to determine whether the cell type in question engulfs synapses in a heterogeneous or a homogeneous manner. (3) The original engulfment assay was optimized for the detection of synaptic material within microglia, rendering the protocol suboptimal for quantifying engulfment by OPCs, which have different morphological structures and make *bona fide* synapses with neurons.

Our protocol addresses these challenges through the use of ratiometric pH-sensors expressed *in vivo.* These sensors can increase the confidence that a synaptic input has been engulfed by an OPC. In addition, instead of the classically applied masking-based approach for quantifying synapse engulfment in Imaris, we have adopted a distance-filtering method based on the optical resolution of the acquisition system. This new filtering-based method is a more rigorous and conservative strategy for classifying synaptic material as engulfed by OPCs (Figure 4a-bii). We experimentally validated this approach as being significantly more conservative in identifying engulfed inputs than the masking-based approach used in the classical engulfment assay (Figure 4c). Furthermore, this protocol extends beyond imaging-based methods by including the flow cytometric analysis of presynaptic proteins within tens of thousands of individual OPCs in each experiment. While other groups have begun to utilize flow cytometry to study synapse engulfment by microglia^18^, the approach has not yet been widely used, standardized, or applied to OPCs (apart from our study^9^). Thus, our protocol provides numerous advantages over other assays, including its specialization for the study of synapse engulfment by OPCs, which have emerged as remarkably multi-functional cells in the brain^9–11,19–21^.

**Figure 4.**
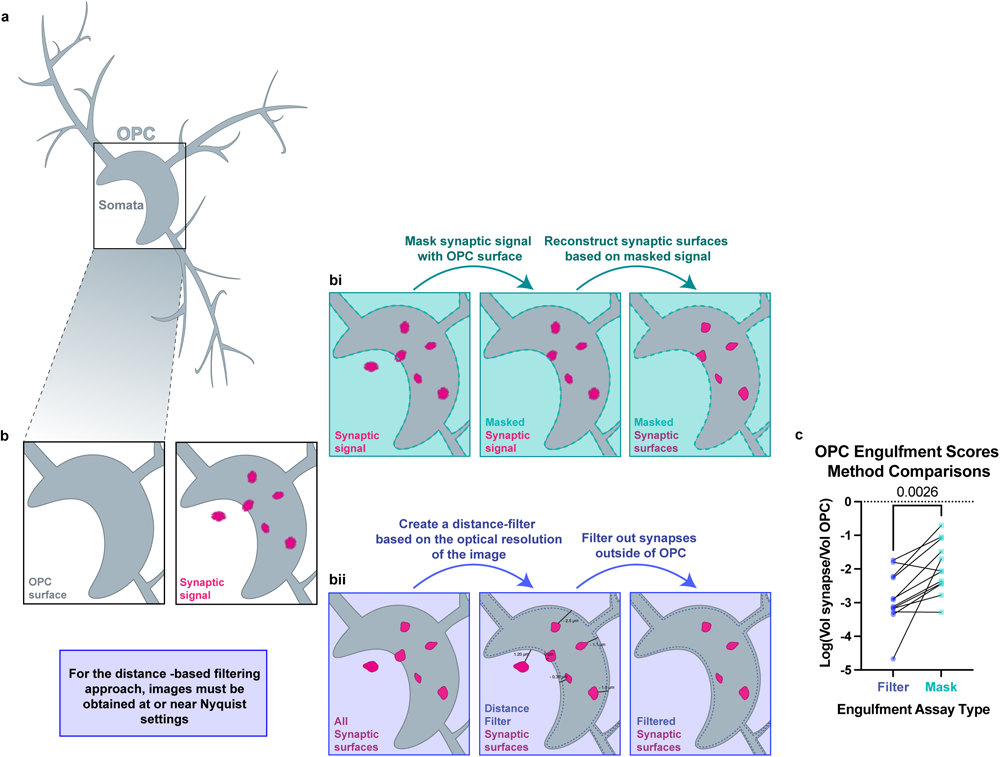
Comparison between engulfment assays performed in Imaris: the traditional masking method versus distance-based filtering method. (a) Confocal images are imported into Imaris in which an OPC surface is reconstructed (grey). (b) The soma of the OPC is visualized along with VGLUT2 signal (magenta) representing presynaptic inputs. (bi) In the traditional method, the OPC surface (teal, dashed line) is used to the mask the VGLUT2 signal, and this masked signal is used to reconstruct the VGLUT2 surfaces (magenta). The volume of the synaptic surfaces contained within the OPC is normalized to the OPC volume to obtain an engulfment score. (bii) Our distance-based filtering approach first creates VGLUT2 surfaces (magenta). Then the optical resolution of the image is used to set an upper threshold for a distance filter (purple dotted line), which defines what is considered ‘inside’ the OPC. Imaris then calculates the distance between the VGLUT2 surfaces and the OPC, and then filters out any VGLUT2 surface not within a certain distance of the OPC surface. The resulting surfaces are used to calculate an engulfment score as described above. (c) The engulfment scores from a single dataset of OPCs (n=12) were calculated using the two different approaches. The distance-based filtering method (purple) resulted in significantly smaller engulfment scores compared to the traditional masking approach (teal), indicating that the distance-based filtering method is more conservative than the masking method. Paired t-test, p = 0.0026.

### Expertise needed to implement the protocol

These approaches combine numerous methods that are relatively standard in molecular and cellular biology laboratories. In particular, the surgical injections of viral constructs should be conducted only by investigators trained to perform these procedures according to institutional policies, and only investigators trained to handle biohazardous agents like AAVs should perform the infections. In addition, flow cytometry experiments require expertise in using fluorescent sorting machinery effectively. The availability of a flow cytometry core facility with trained staff could circumvent this requirement.

### Limitations of the approach

While the approaches that we describe here represent a series of advancements compared to other methods (as described above), it is important to acknowledge the limitations of these strategies. One limitation of this strategy is that each of these approaches relies upon the use of antibodies to detect specific proteins. Given that antibodies require rigorous validation and optimization and are subject to manufacturing variability between lots, integrating new antibodies into the protocols can add more time to the front-end of experiments for an investigator who wants to adapt the approach, for example to quantify synapse engulfment by microglia rather than OPCs. Another limitation is that our suggested distance filter-based engulfment analysis in Imaris results in the loss of information in thinner regions of the cell, which can include distal processes. Thus, our method focuses on larger regions, which primarily include OPC somata and proximal processes. Regarding the flow cytometry strategy, it is important to note that structural parameters, such as whether an input resides within the processes or cell body of an OPC, are lost due to tissue homogenization. Finally, another limitation is that these approaches are optimized for the rodent brain and adapting them to other models would likely require substantial adjustments. Despite these caveats, we expect the approaches described here to significantly increase the rigor and reproducibility of experiments quantifying synapse engulfment by OPCs and other glial cell types as well.

### Experimental design

This protocol describes two complementary methodologies to analyze the engulfment of synaptic material by OPCs. In the imaging-based approach, OPC engulfment of thalamocortical synapses in primary visual cortex is described. However, these protocols can be adapted for different brain regions and can be used to compare OPC engulfment across conditions and in disease models. It is suggested that experimenters engage in robust validation of the methods when used in different contexts. For example, we recommend validating new injection coordinates in the mouse brain for investigators who are interested in quantifying engulfment in non-visual regions. It is also important to optimize antibody staining in the different regions as well.

The experimental design for flow cytometry can be complex due to the large number of controls that are required to prepare for cell-specific target protein analysis. We designed the fluorophore panel using the FluoroFinder platform, however many other manufacturer websites can help to accomplish this task. When choosing the fluorophore, assign the brightest fluorophore to the lowest expressed protein and the dimmest fluorophore to the highest expressed protein. Avoid fluorophores with emissions that have high spectral overlap and always include the appropriate controls for the experiment as follows. Every experiment should include an unstained control to avoid autofluorescence, single color controls for compensation of spectral overlap, viability control to discriminate live cells from dead cells and fluorescence minus one (FMO) for a reliable gating strategy (Figure 5). CRITICAL In order to choose the correct fluorophore to be used in the experiment, it is necessary to know which lasers and optical filters are available on your system. In the present manuscript, three experimental tubes and six control tubes were prepared, including two control tubes containing cells (Unstained control and FMO-SYN1) and four control tubes containing beads for single stained control (Figure 3a). CRITICAL To avoid systematic errors, label the control and experimental tubes with different colors, and during the procedure, separate the tubes that will be used for cells from the tubes that will be used for beads.

**Figure 5.**
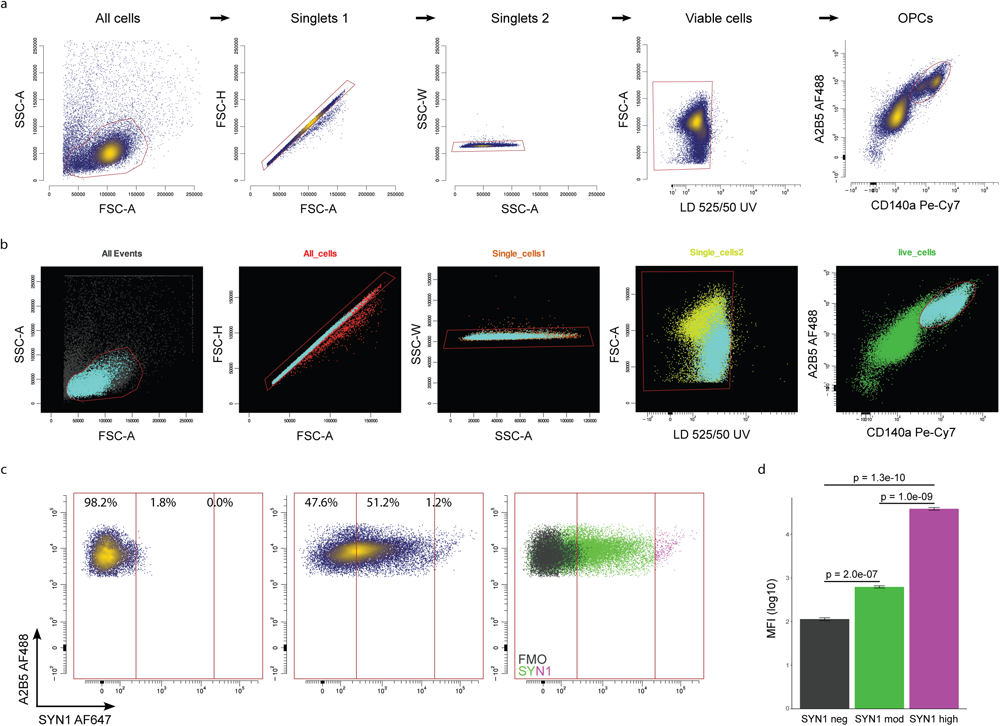
Gating strategy for identifying oligodendrocyte precursor cells (OPCs) and measuring the intracellular content of SYN1. (a) First, cells of interest are identified based on size and granularity using a side scatter (SSC-A) vs. a forward scatter (FSC-A) plot. Doublet events are excluded by plotting FSC-H versus FSC-A (Singlets 1) followed by SSC-W versus SSC-A (Singlets 2). Dead cells and debris are then removed using a LIVE/DEAD aqua stain. The OPC population is defined based on the specific markers A2B5 and CD140a (PDGFRA). (b) Back-gating is applied to ensure the correct gating strategy for OPC identification between different samples. (c) Representative flow cytometry plots show the variable intracellular content of SYN1 in OPC populations (green, OPCs containing a moderate amount of SYN1; magenta, OPCs containing a large amount of SYN1). A fluorescence minus one (FMO) control (grey) is used to set the gate for the negative and positive populations. (d) A bar plot showing the mean fluorescence intensity (MFI) of SYN1 in the different OPC populations. The highest MFI in the OPC population is used to set the gate for the heaviest engulfers (magenta in [c]). SYN1 neg, OPCs not containing synaptic material; SYN1 mod, OPCs containing a moderate amount of synaptic material; SYN1 high, OPCs containing a large amount of synaptic material. Statistical analysis: Pairwise t-test followed by Bonferroni multiple test correction. SYN1 mod vs SYN1 neg, p = 2.0e-07; SYN1 high vs SYN1 neg, p = 1.3e-10; SYN1 high vs SYN1 mod, p = 1.0e-09.

## SECTION A – Imaging-based protocol

### Materials

#### Animals

To encourage reproducibility, animals should be from the same background strain of C57Bl/6J. The approach will likely be adaptable to multiple mouse lines, but we have not yet performed experiments in other strains. Animals used for immunofluorescent staining were wild-type C57Bl/6 mice (Jackson Laboratory strain code B57BL/6J).

▴CAUTION All animal husbandry and experiments were conducted following the policies established by the Institutional Animal Care and Use Committee (IACUC) at Cold Spring Harbor Laboratory. All experiments were previously approved by the IACUC.

### Reagents

#### Reagents for animal perfusion and tissue fixation

- Gibco 1X phosphate-buffered saline (PBS), pH 7.4 (FisherScientific cat. no. 10010049)
- Paraformaldehyde 16% Aqueous Sol. EM Grade (Electron Microscopy Science cat. no. 15710)
- D-(+)-Sucrose (VWR cat. no. BDH9308-500G)
- Dry ice

#### Reagents for immunofluorescence staining

- Gibco 1X phosphate-buffered saline (PBS), pH 7.4 (FisherScientific cat. no. 10010049)
- TRITON™ X-100 (VWR cat. no. 97063-864)
- Gibco™ Fetal Bovine Serum (Life Technologies cat. no. A3160501)
- Normal Goat Serum (Thermo Fisher cat. no. 31873)
- Normal Donkey Serum (Jackson ImmunoResearch cat. no. 017-000-121)
- Primary Antibodies (Table 1)

#### Reagents for surgery

- pAAV:hSYN-synaptophysin-mCherry-eGFP (pSynDig)
- Betadine (Amazon cat. no. B005R8580M)
- Metacam (Boehringer-Ingleheim cat. no. 136327)
- Isoflurane (MWI Veterinary Supply cat. no. 502017)
- Buprenorphine HCl Inj. (Covetrus cat. no. 059122)
- Gibco 1X phosphate-buffered saline (PBS), pH 7.4 (FisherScientific cat. no. 10010049)
- Flunixin Meglumine (Covetrus cat. No. 11695-4025-1)

#### Equipment General equipment

- Watson Marlow 205CA4 Channel pump with Pump Pro MPL (Boston Laboratory

Equipment cat. no. BLE2000180) CRITICAL This system can be replaced with other perfusion pumps.

- PVC Tubing (1/16 x 1/8 in) (Sigma-Aldrich cat. no. Z280348)
- 22-gauge needle with tip cut off (VWR cat. no. BD305156)
- Nalgene® desiccator (Sigma-Aldrich cat. no. D2672-1EA)
- Mini Dissecting Scissors, 8.5c (World Precision Instruments [WPI] cat. no. 503667)
- Operating Scissors straight 11.5 cm (WPI cat. no. 501753)
- Dumont Tweezers #5 (WPI cat. no. 501985)
- Dressing Forceps (WPI cat. no. 500363)
- Kimberly-Clark Professional™ Kimtech Science™ Kimwipes™ Delicate Task Wipers (Fisher Scientific cat. no. 06-666A)
- Embedding molds (VWR cat. no. 15160-215)
- Tissue-Tek® O.C.T. Compound, Sakura® Finetek (VWR cat. no. 25608-930)
- Aluminum Foil (VWR cat. no. 89107-724)
- 15 mL Falcon® Centrifuge Tubes, Conical Bottom (VWR cat. no. 21008-918)
- 50 mL Falcon® Centrifuge Tubes, Conical Bottom (VWR cat. no. 21008-951)
- Corning® bottle-top vacuum filter system (Sigma Aldrich cat. no. CLS431205)
- Leica Microsystems 3P Glass Insert 70 mm Wide for Anti-Roll Systems (Fisher Scientific cat. no. NC0470572)
- 30 mm Specimen Chuck--green O-ring (VWR cat no. 10756-204)
- 200 Proof KOPTEC Ethanol (VWR cat. no. 89125-186)
- Fisherbrand™ Superfrost™ Plus Microscope Slides (Fisher Scientific cat. no. 12-550-15)
- VWR® Microscope Slide Boxes for 100 Slides (VWR cat. no. 82003-406)
- Surgipath® Low-Profile 819 Disposable Sectioning Blades (VWR cat. no. 10015-014)
- Razor blades (VWR cat. no. 55411-050)
- Paint brushes (Amazon cat. no. B07GH7WGC3)
- Gloves (Fisher Scientific cat. no. 19166096)
- NitroTAPE Cryogenic Tape (Thomas Scientific cat. no. 1184W61)
- HybEZ™ II Hybridization System for Manual Assays (ACD cat. no. 321710-R)
- ImmEdge™ Hydrophobic Barrier Pen, Vector Laboratories (VWR cat. no. 101098-065)
- DAPI Fluoromount-G® (Southern Biotech cat. no. 0100-20)
- Fluoromount-G® (Southern Biotech cat. no. 0100-01)
- Fisherbrand™ Cover Glasses: Rectangles (Fisher Scientific cat. no. 1254418P)
- Cryostat Leica CM3050S
- Zeiss Confocal Laser Scanning Microscopy (LSM) 710 or LSM 780 microscope with ×20/0.8 NA (air) and ×63/1.4 NA (oil) objectives

**Table 1:** Overview of the antibodies used in these protocols.

| Antibody | Host animal | Marker | Cat # | Usage | Dilution |
| --- | --- | --- | --- | --- | --- |
| VGLUT2 | guinea pig | Thalamocortical excitatory projections | AB2251-I | IF | 1:1000 |
| NG2 | rat | OPC | MA5-24247 | IF | 1:250 |
| MBP | rat | Myelin/oligodendrocytes | AB7349 | IF | 1:1000 |
| Sox10 | rabbit | OL lineage | AB227680 | IF | 1:100 |
| Lamp2 | rat | Lysosome | AB13524 | IF | 1:200 |
| PDGFRA | goat | OPC | AF1062 | IF | 1:500 |
| A2B5 Alexa Fluor 488 | mouse | OPC | FAB1416G | FC | 1:100 |
| CD140a PE-Cy7 | rat | OPC | 135912 | FC | 1:100 |
| SYNAPSIN-1 Alexa Fluor 647 | rabbit | Synaptic material | 111275 | FC | 1:100 |
| CD16/CD32 | rat | blocking antibody | 14-0161-82 | FC | 1:1000 |

**Table 2:**
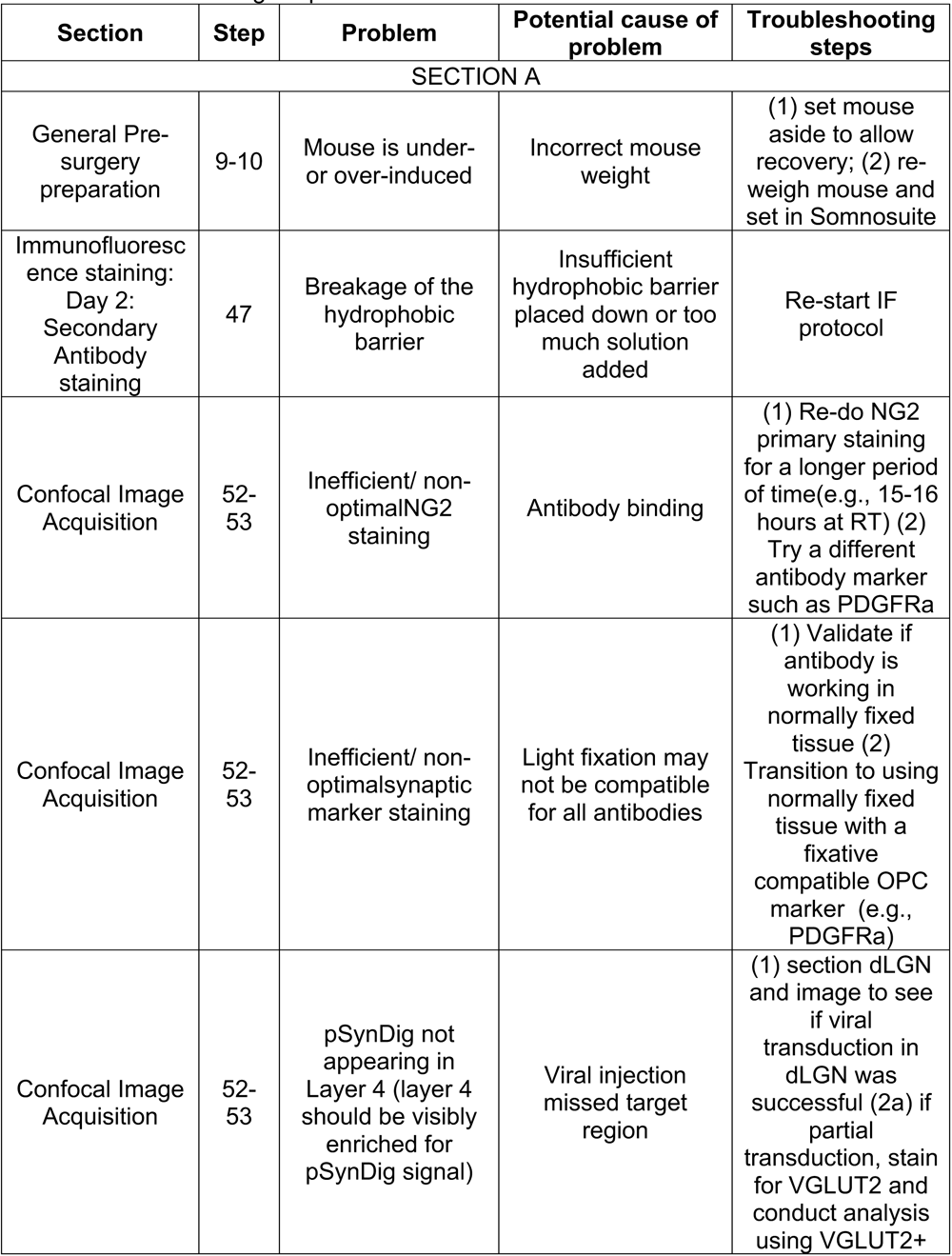
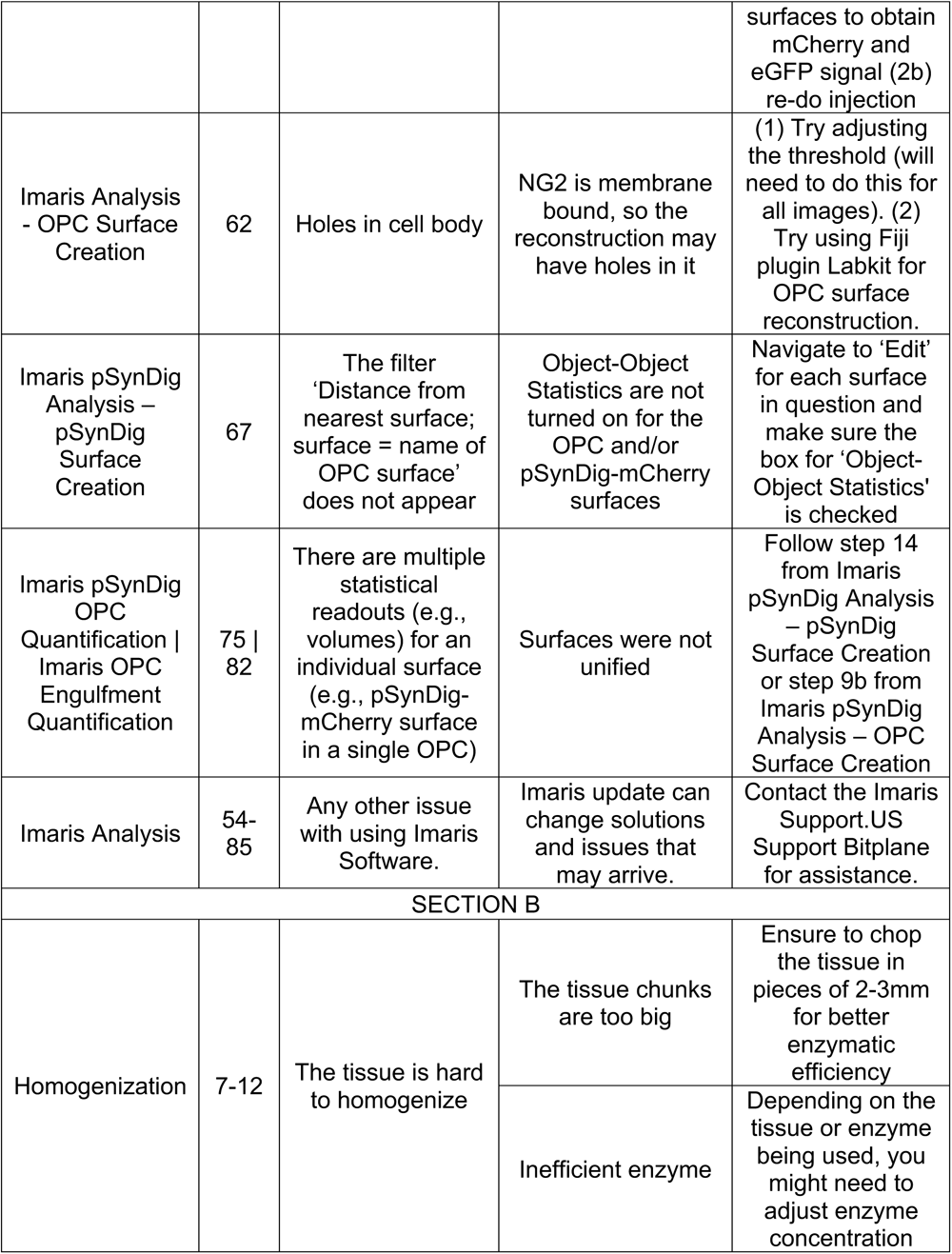
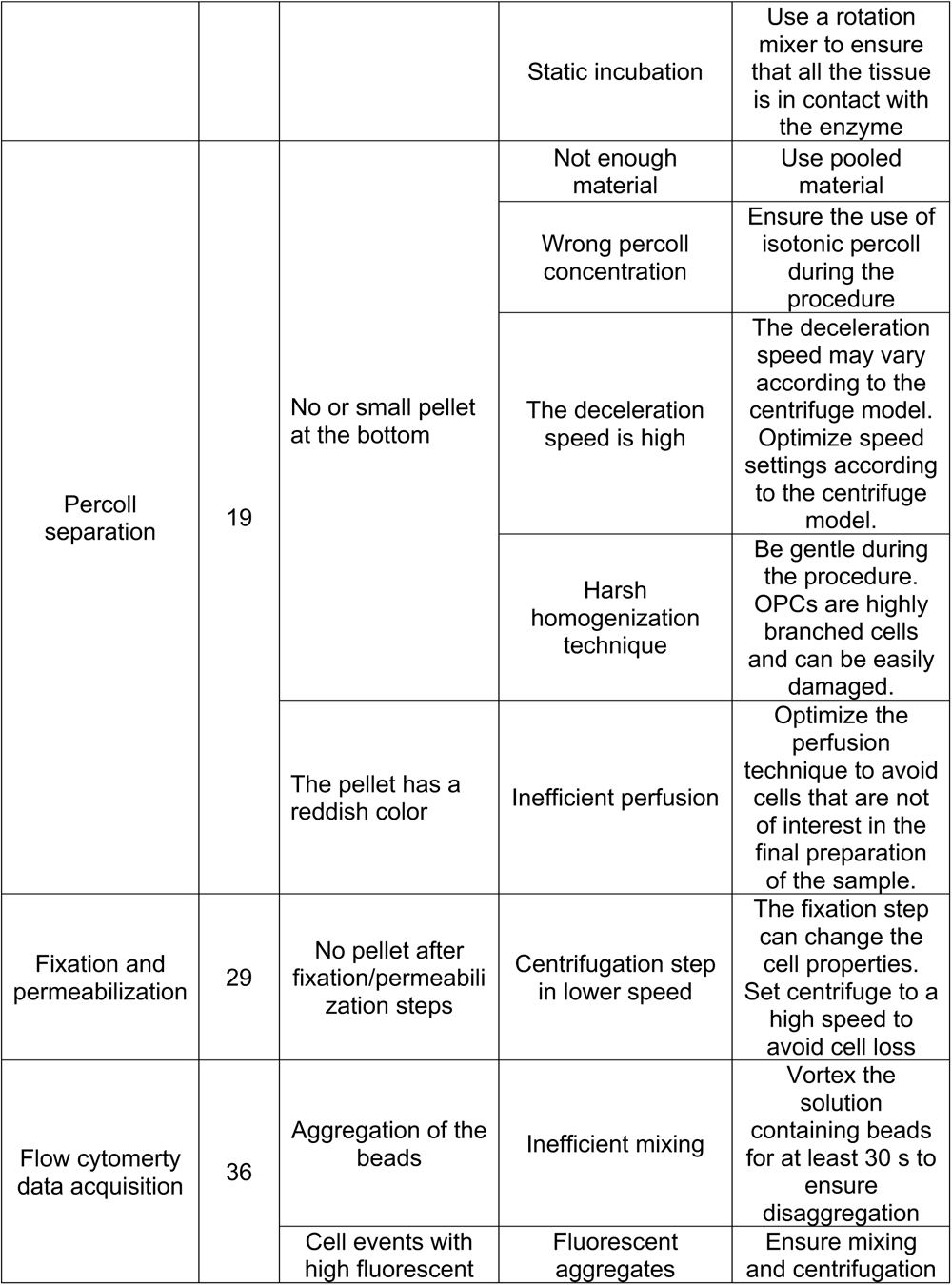
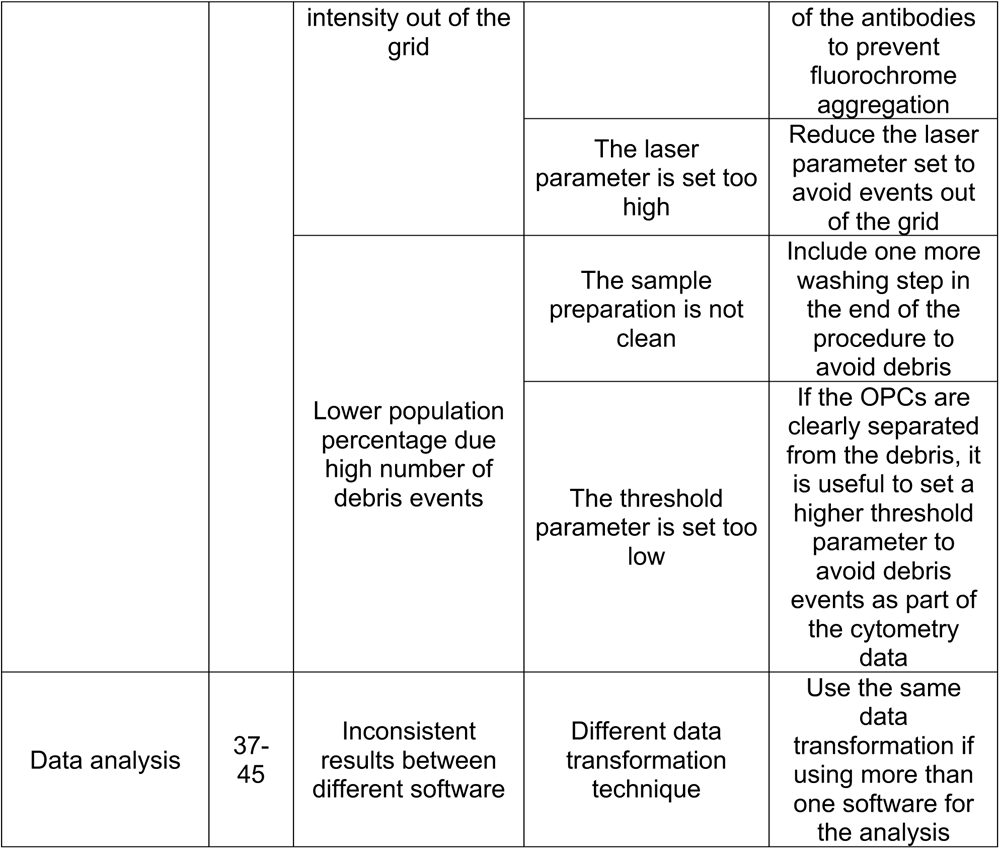
Troubleshooting Steps.

#### Surgery Equipment

- 69100 Rotational Digital Stereotaxic Frame for Mice and Rat (RWD cat. no. 69100)
- Stereotaxic Frame Nosecone Masks (RWD cat. no. 68601)
- Leica M50 Stereomicroscope with Leica M50 optics carrier (Leica cat. no. 10450154) AND SMS 25 Articulating Arm Stand with Focus Mount and 90 Degree Adapter (Leica cat. no. 8096629)
- Ear bars (Stoelting cat. no. 51649)
- Somnosuite (Kent Scientific cat. no. SS-01)
- Somnosuite starter kit (Kent Scientific cat. no. SOMNO-MSEKIT)
- Activated charcoal filters (Kent Scientific cat. no. 10-2001-8)
- Heating Pad (Amazon cat. no. B018VQ72RI)
- Scalpel blades (Harvard Apparatus cat. no. 75-0088)
- VetBond (Amazon cat. no. B079QJXK46)
- Scalpel Handle (Harvard Apparatus cat. no. 72-8686)
- Scalpel blades (Harvard Apparatus cat. no. 75-0093)
- Microdrill (RWD cat. no. 78001)
- 0.45 mm Drill bits (Stoelting cat. no. 514551)
- Motorized injector (Stoelting cat. no. 53311)
- 34-gauge, Small Hub RN Needle (Hamilton; custom) (Hamilton Company cat. no. 207434)
- Neuro Syringe (Hamilton Company cat. no. 65460-03)
- Gelfoam (VWR cat. no. 10611-588)
- Dry bead sterilizer (Kent Scientific cat. no. INS700860)
- MAXI CARE™ Underpads, Covidien (VWR cat. no. 82004-836)
- Surgical Gloves (VWR cat. no. 89411-648)
- Sterilization trays (VWR cat. no. 100498-918)
- Sterile cotton tip applicator 3” (VWR cat. no. 76407-736)
- Sterile cotton tip applicator 6” (VWR cat. no. 76407-738)
- VWR® Syringe Filters (VWR cat. no. 28145-501)
- Insulin syringes/Tuberculin syringes 1CC 26GX3 (Fisher Scientific cat. no. 14-823-2E)
- Eye lubricant/ Genteal Tears Ophthalmic Gel (Covetrus cat. no. 72359)
- Nair Hair Remover Lotion Aloe & Lanolin 9oz (Amazon cat. no. B078YGW7Q3)

#### Data analysis software

- Fiji^22^ (ImageJ2, v.1.51., https://fiji.sc/)
- Imaris (v.10.0., Andor)
- Imaris File Converter (v.10.0., Andor)
- GraphPad Prism (v.9.4. for Mac, GraphPad Software, San Diego, California USA, www.graphpad.com)
- Microsoft Excel (v.16.70., Microsoft)
- Zen black 2012 SP5 (v.14.0. Zeiss) for acquisition from LSM710
- Zen black 2011 SP7 (v.14.0. Zeiss) for acquisition from LSM780

#### Reagent setup 4% (vol/vol) PFA

Dilute 16% PFA with 1X PBS to obtain 4% PFA. Solution can be stored for a few weeks at 4°C.

▴CAUTION Toxic reagent. Always handle with gloves and avoid eye and skin contact. Handle reagent in hood with high airflow. Only use with designated PFA tools.

### 15% and 30% (wt/vol) sucrose solution

Weigh out the appropriate grammage of sucrose and dissolve in 1X PBS to obtain a 15% and/or 30% (wt/vol) sucrose solution. Filter this solution through a bottle-top vacuum 0.22 µm filter system and store long-term at 4°C.

### 10% TritonX-100

Dilute TritonX-100 with 1X PBS until a 10% TritonX-100 solution is achieved. Cover in aluminum foil and place on a rocker at RT until dissolved. Store at 4°C long-term covered in aluminum foil.

### Blocking Solution

Use the 10% TritonX-100 solution to create a solution with a final concentration of 0.3% TritonX-100 in 1X PBS. To this same solution, add an appropriate volume of Normal Goat Serum (NGS), Fetal Bovine Serum (FBS), or Normal Donkey Serum (NDS) to reach a final concentration of 5% NGS/FBS/NDS. The type of serum used should be optimized according to the desired secondary antibodies (e.g., if secondary antibodies with a Donkey host are used, use a NDS-based blocking solution).

### Probing Solution

Use the 10% TritonX-100 solution to create a solution with a final concentration of 0.1% TritonX-100 in 1X PBS. To this same solution, add an appropriate volume of NGS, FBS, or NDS to reach a final concentration of 5% NGS/FBS/NDS. The probing solution should contain the same serum as the blocking solution for a given experiment.

### TritonX-100 in PBS (PBST)

Use the 10% TritonX-100 solution to create a solution with a final concentration of 0.1% TritonX-100 in 1X PBS.

### 70% (vol/vol) Ethanol

Dilute 200 Proof (100%) Ethanol with MilliQ water until a 70% Ethanol solution is achieved. Store at room temperature for long-term storage.

▴CAUTION Flammable reagent. Keep in appropriate storage with other flammable reagents and keep away from open flames.

#### Flunixin meglumine (0.5 µg/mL)

Dilute stock Flunixin meglumine in sterile saline solution (0.9%). Store at room temperature until the expiration date.

#### Equipment setup SURGERY SETUP

We assembled the surgical set up, including stereotaxic apparatus with Somnosuite isoflurane delivery and stereomicroscope, based off the manufacturers’ guidelines.

## Procedure

### General pre-surgery preparation ●30 minute

<u>Note</u>: All surgical tools should be sterilized via autoclave prior to surgery.

▴CAUTION Surgical procedures should be conducted in accordance with the suggested procedures set by the institution. Steps 1-25 should be adapted based on institutional guidelines.

▴CAUTION This surgery involves usage of an adeno-associated virus (AAV) and all procedures should comply with Institutional Biosafety Committee (IBC) guidelines and policies regarding usage of AAVs.

1. Put on surgical gloves, mask, and appropriate personal protective equipment.
2. Pre-warm heating pad mounted on the stereotaxic frame stage to 37°C.
3. Place a sterile surgical drape over the heating pad to create a sterile field. Place autoclaved and sterile surgical tools in sterilizer tray when not in use.

a. Use the hot bead sterilizer to sterilize any tools that touch nonsterile areas.
4. Weigh animals and calculate in advance the volumes of drugs which will be administered in later stages of the surgery.
5. Provide mice with an S.Q. injection to Meloxicam [10 mg/kg] at least 3 hours prior to the surgery.
6. Insert the weight of the mouse into the Somnosuite for appropriate isoflurane delivery.
7. Anesthetize the mouse by placing it into the induction chamber infused with vaporized isoflurane at an **initial flow rate of 3-4%.** Ensure the isoflurane lines are directed to the induction chamber and not the nose cone on the stereotaxic apparatus.
8. Once the mouse is deeply anesthetized, place the mouse onto the pre-warmed surgical set-up/stereotaxic frame.
9. **Adjust the flow rate to 1.5-2%** to maintain the deep anesthetization and redirect the flow of isoflurane to the stereotaxic apparatus. Slide the mouse into the nosecone and fasten the nosecone into place. TROUBLESHOOTING
10. CRITICAL STEP Conduct toe/tail pinch test to confirm anesthetic depth and look for decreased respiratory rate. If the mouse fails the toe/tail pinch or has a decreased respiratory rate, adjust the flow rate of isoflurane. DO NOT EXCEED 2% flow rate. Ensure that respiration is steady and controlled and if respiration includes lurching decrease isoflurane flow rate. DO NOT GO BELOW 1.5%.
11. Place mouse into ear bars and lock mouse in place.
12. CRITICAL STEP Apply a liberal amount of eye lube onto a sterile cotton swab and roll the lubricant onto the mouse’s eyes to prevent them from drying out.
13. Remove hair from the surgical site by applying Nair using a cotton swap onto the scalp of the mouse in circular motions. Leave the Nair on for about 30 seconds and remove excess with a cotton swab saturated in de-ionized water or 1X PBS. Repeat this process until the intended surgical site is cleared of hair. CRITICAL STEP Do not leave Nair on for longer than 30 seconds as you will increase the chance of chemical burns, which are highly irritating to the mouse. Make sure all Nair is removed prior to proceeding.
14. Apply an antiseptic, bactericide Betadine soap (Betadine scrub, containing povidone iodine) to the surgical site using a sterile cotton swab. Start at the incision site and work outwards. Clean with a cotton swab saturated with 70% ethanol following the same motions. Repeat this betadine-ethanol application a total of three times.

### Stereotaxic Injection ● 1-2h

15. Apply topical **bupivacaine (.75%)** to the surgical site.

b. Corticosteroids (**dexamethasone 0.5 mg/kg IP or methylprednisolone 30 mg/kg**) may also be administered through IP injections at this time to minimize both intraoperative brain swelling and post-operative inflammation and gliosis around the injection site.
16. Prior to starting, again check the respiration rate and anesthetic depth of the mouse and adjust the isoflurane flow rate as needed. Do not exceed the range of 1.5-2% flow rate.
17. Using a new, sterile scalpel blade make an anterior to posterior incision along the scalp of the mouse. Retract the skin and expose the surface of the skull.
18. Use a sterile cotton swab applicator to push back and clear the area of the periosteum. Resolve any bleeding at the surface immediately with sterile cotton swab.
19. Attach a sterilized, clean Hamilton syringe to the microinjector onto the swinging arm of the stereotaxic apparatus.

c. Gently lower the needle into an aliquot of pSynDig virus and aspirate 1.2 – 2X the volume of virus intended to be injected. Ensure there are no clogs by infusing a small amount of virus back into the aliquot. ▴CAUTION Handling of AAVs should be conducted in accordance with institutional policies and be handled in a BS2 approved facility.

d. Lock the swinging arm of the stereotaxic apparatus into place such that the arm is set to 0 degrees in relation to the base of the apparatus.
20. Using the adjustment knobs on the stereotaxic frame, adjust the mouse skull so Bregma and Lambda are within 0.05 mm in X, Y and Z coordinates (relative to Bregma’s position). Further adjust the roll of the mouse’s skull to ensure that the skull is flat and within an error of 0.05 mm.
21. Once level, use Bregma to identify the location of the insertion site for needle (adult coordinates for the dLGN are X: ±2.15, Y: -2.15 according to Bregma) on one or both sides of the brain.

e. Use a sterilized microdrill with a 0.45 mm drill tip to drill a small entry hole (approximately the size of the drill bit) into the skull of the mouse at the injection site. Make sure not to go deep enough to injure the mouse brain.
f. Use gel foam soaked in filtered, sterile 1X PBS to resolve any bleeding that occurs.
22. Conduct stereotaxic injection of pAAV:hSYN-synaptophysin-mCherry-eGFP (pSynDig) into dLGN of the thalamus.

g. Lower the needle until you reach the brain tissue and zero the Z position. SLOWLY enter the brain tissue at a rate of ∼0.01 mm/3 seconds.
h. Go 0.05 mm lower than the Z coordinate and pause for 3 minutes. Raise the needle to the correct Z coordinate (adult dLGN Z-coordinate: -2.9).
i. Set the volume and rate of injection to 50 nL/minute. Hit ‘Start’ on the motorized injector to begin the injection.

Inject 250 nL of each virus (5×10^12^ titer) into each dLGN.
j. After the injection finishes, wait for 10 minutes. After the wait, SLOWLY remove needle at a similar rate to when entering the brain.
23. Repeat steps 21-22, if desired, with the contralateral hemisphere.
24. Place a few drops of sterile 1X PBS onto the incision site to help loosen skin. Bring skin together and seal with small quantity of vetbond (use minimally to avoid irritation).
25. Inject the mouse with Flunixin meglumine [2.5 mg/kg; i.p.]. Place the animal in a separate cage on a heating pad apart from the others to let it recover before placing it back with other animals. Make sure the mouse always has access to food and water (hydrogel). CRITICAL STEP Wait for the mouse to become active again (∼15-45 minutes) before placing back into colony. Fill out appropriate post-surgery documentation required by the animal services at your institution.

### Post-surgery ● 2-3 weeks

26. Check and monitor the mouse’s condition twice daily and follow all institutional policies for post-surgical animal welfare.

k. Apply topical bupivicaine (0.75%) as needed after surgery not to exceed one application every 24 hours.
l. Administer Meloxicam once daily on the day of surgery (see pre-surgical prep) and then as needed.
27. PAUSE POINT Wait at minimum 2-3 weeks after surgery to permit the virus to adequately express. After this period, proceed onto Animal sacrifice and perfusion.

### Animal sacrifice and perfusion for immunofluorescence staining ● 2-3 d

28. Euthanize the animal with a method appropriate for perfusion. We suggest using isoflurane by placing the mouse into a Nalgene desiccator containing a Kimwipe saturated with about 200 µL of isoflurane. Wait until the mouse is no longer responsive and remove from chamber. Maintain isoflurane anesthesia and conduct strong toe and tail pinches to ensure animal is deeply anesthetized prior to proceeding.

### Processing samples for anti-NG2 antibody staining to label OPCs

29. Connect a 22-gauge needle to a Watson Marlow 205CA4 Channel pump with Pump Pro MPL pump through a line of PVC tubing and insert needle into the left ventricle of the heart. Set the pump rate to 30 rpm and perfuse approximately 10 mL of ice cold 1X PBS or until the liver clears.
30. After perfusion, extract the brain from the mouse and place into 10 mL of cold 4% PFA in 1X PBS in a labeled 15 mL falcon tube. As soon as the mouse brain touches PFA start a 2-hour timer.

CRITICAL STEP Drop-fix the brain for 2 hours at 4°C. If brain is fixed for longer, NG2 staining will not work well.

31. Wash the brain 3X in cold 1X PBS.
32. Place the washed brain into a 30% sucrose solution and allow the brain to sink overnight.

### Embedding, freezing, and storing samples

33. Embed brain in OCT using embedding molds. Make sure brain is straight, level, and completely submerged in OCT. Place onto flat plane of a dry ice slab and cover with aluminum foil until frozen.

a. PAUSE POINT Store at −20°C temporarily prior to sectioning and move to −80° C for long term storage.

### Cryosectioning ● 3-5 h

34. Obtain embedded, frozen brain from −20°C storage. Place the following items into the −20°C cryostat chamber to permit objects to come to temperature (brain, chuck, anti-roll plate, 819 blade, razor blade and paint brushes).
35. Set Objective temperature to −17°C and chamber temperature to −21/20°C, move objective all the way back and adjust the stage base angle to 5°, and adjust section thickness to 25 µm.
36. Attach the embedded section to the chuck using OCT. Freeze the brain to the chuck with anterior side of the brain facing the chuck. Section from the posterior side of the brain to reach the visual cortex sooner.
37. Place the chuck-mounted section into the microtome specimen holder and begin trimming the excess OCT in 50-100 µm intervals until the brain is exposed. Make sure to adjust the angle of the specimen using the adjustment lock to obtain symmetric brain sections. Continue trimming and discarding trimmed sections until desired brain region is approached (in this case, visual cortex).
38. Turn off trimming function to start obtaining 25 µm sections. Place the anti-roll plate down onto the stage and begin collecting sections. Mount sections directly onto the slide. Collect 2-4 sections onto each slide until the visual cortex has been sectioned completely.
39. PAUSE POINT Store slides with brain sections in a slide box at −20°C for a few months prior to immunostaining. Move to −80°C for long term storage.

### Immunofluorescence staining ● 2 d Day 1: Primary Antibody staining

40. Create a dark-humidity chamber by placing damp paper towels in a box (e.g., Slide box) and place sections inside, lying flat.
41. Wash sections in 1X PBS for 5-10 minutes to remove OCT. Gently dry the slide with a kimwipe and make sure all the OCT and PBS is removed.
42. **Optional but recommended:** Bake slides for 15-30 mins at 60°C using a HybEZ™ II Hybridization System or another bench-top oven. This will help prevent the tissue from lifting throughout the staining process.
43. Place dried slides back into the staining chamber and wash sections in 200 µL per section of PBST (0.1% TritonX-100) for 5-10 minutes.
44. Draw 2-3 concentric hydrophobic barriers around each section with the ImmEdge™ Hydrophobic Barrier Pen. Allow the barriers to dry for 10 min or until visibly dry.
45. Block the samples with 100 µL of blocking solution (see Reagents setup) for 1 hour at room temperature in the dark. This volume may need to be adjusted to ensure that a sufficient volume of blocking solution is submerging the entirety of the tissue sample.
46. Prepare 100 µL per sample of primary antibody in the probing solution (see Reagents setup) by adding the appropriate primary antibodies into the solution at desired concentrations (Table 1). CRITICAL STEP The antibody concentrations in Table 1 have been optimized through antibody titration trials. These concentrations may need to be further optimized for independent experiments conducted in other laboratories.

a. Remove the blocking solution and replace it with the primary antibody solution. Incubate sections in primary antibody solution either overnight at 4°C or at room temperature for 1 hour.

### Day 2: Secondary Antibody

47. Remove the primary antibody solution from sections and wash 3X in PBST for 10 minutes. TROUBLESHOOTING
48. Prepare the secondary antibody solution in the same probing solution as for the primary antibody. Incubate in secondary antibody solution for 1 hour at room temperature.
49. Wash 3X with 100 µL per sample 1X PBS for 10 minutes.
50. Add 20 µL of Fluoromount-G with or without DAPI (depending on staining scheme) per sample and coverslip.
51. PAUSE POINT Store at 4°C protected from light.

### Confocal Image Acquisition ● 3 to 9 h

52. Acquire confocal images using a LSM 710 or LSM 780 (Zeiss) microscope with either a ×40/1.3 NA (oil) or ×63/1.4 NA (oil) objective. Use the 63X objective to acquire images containing 1-3 OPCs. TROUBLESHOOTING

CRITICAL STEP Image at Nyquist settings for optimal data acquisition.

53. These settings are retained throughout the imaging of the entire experiment across all conditions to increase reproducibility. Volumetric Z-stacks were acquired to capture the majority of the OPC somata within layer 4 of visual cortex. PAUSE POINT Store data for later analysis.

### Imaris Analysis ● 6 h to 4 d File Conversion

54. The LSM 710 or LSM 780 confocal scopes save files as .czi files. Open the Imaris File Converter application and drag .czi files into the console and select ‘Start All’ and choose appropriate place to save converted files.
55. Files will be converted into .ims files and can then be opened using Imaris software. Open .ims files and organize files according to experiment.

### Image Processing

56. From the Arena, open an image. Navigate to Image Processing by selecting ‘Image Proc’ on the Ribbon (Figure 6a, teal box). Apply the following functions (Figure 6ai) in the given order for the channels of both the OPC and the marker(s):

a. Select ‘Gaussian Blur’ from the drop-down menu then select ‘Ok’.
b. Select ‘Background Subtraction’ from the drop-down menu and select ‘Ok’. Use default settings for both.
57. During Step 3, make sure to confirm that all images have at least one NG2 cell within layer 4 of V1. If not, remove the image from the Arena and do not analyze.
58. Calculate the mean intensity of each of the relevant channels in Fiji.

a. Install the Fiji (ImageJ) plugin in Imaris.
b. Navigate to Fiji through the Imaris Application Taskbar by selecting ‘Fiji’ è

‘Image to Fiji’ (Figure 6av).

i. Once your image is opened in Fiji, apply a Maximum Intensity Projection by selecting ‘Image’ è ‘Stacks’ è ‘Z-project’ (max intensity) for all channels.
ii. In a new window with the Maximum projections of the relevant channels, measure the mean intensity of said channels. Set the measurements by going to ‘Analyze’ è ‘Set Measurements’ and checking the box next to ‘Mean gray value.’ Hit ‘OK’ to apply.

**Figure 6.**
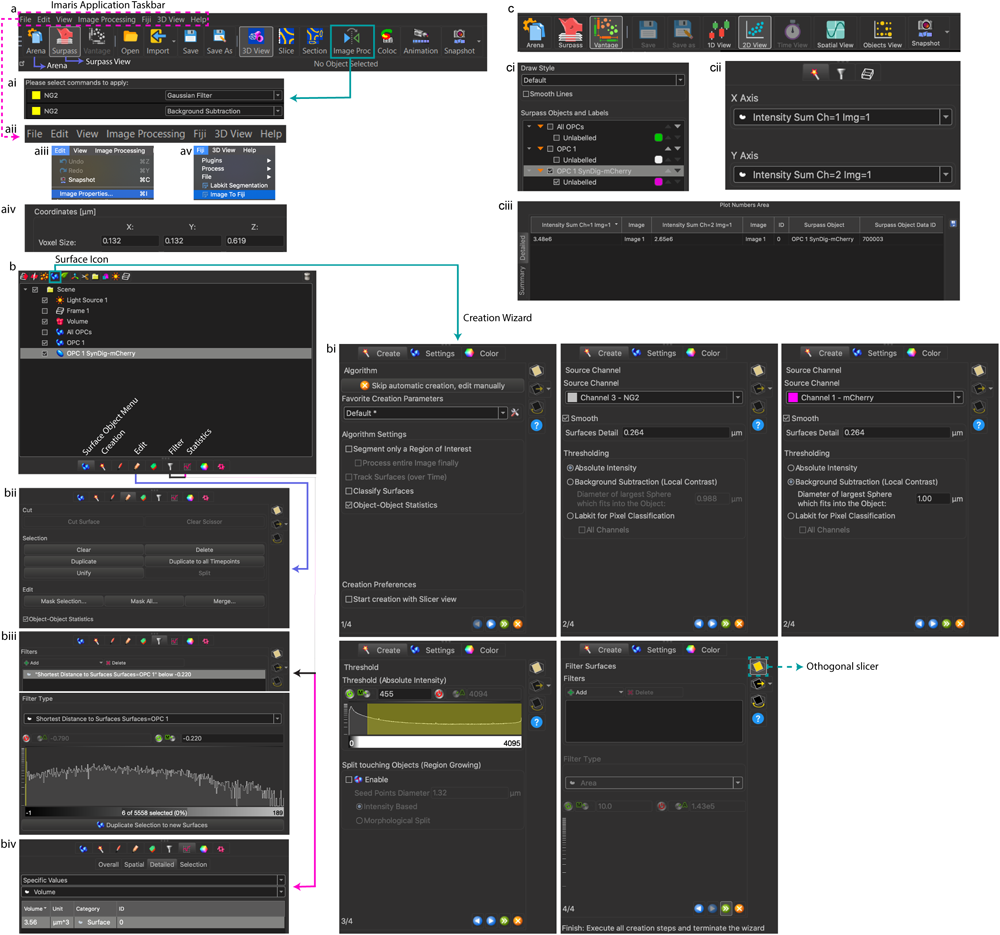
Imaris software workflow. (a) Initial navigation between the Arena and the Surpass View can be done using the respective icons on the ribbon (purple arrows). (ai) Process the images by applying a Gaussian Blur and Background subtraction (teal box and arrow) on relevant channels like NG2, mCherry, and eGFP. (aii) Go to the Imaris Application Taskbar (pink hashed box and arrow) to navigate to (aiii) image properties and record information about (aiv) image resolution which is necessary to calculate threshold values. Mean intensity values can be obtained from (av) the Fiji plug-in. (b) Imaris Creation Menu and Surface Object Menu. Create surfaces by clicking the Surface Icon (teal box). (bi) Following the Creation Wizard as displayed. It is useful to utilize the Orthogonal Slice (teal hashed box) during this process. After surfaces are created, (bii) Unify surfaces under the ‘Edit’ tab (purple arrow), (biii) filter synapse surfaces using the distance filter (black arrow), and (biv) extract volumetric information if desired (pink arrow). (c) pSynDig sum intensity data extraction. Select desired images and navigate to Vantage and 2D View. (ci) Only select surfaces of interest and (cii) set the X and Y axes to obtain the desired Intensity Sum from the eGFP and mCherry channels. (ciii) Navigate to the Plot Numbers Area to check sum intensity data and export using the Save icon. All Imaris Software images were obtained and used with explicit permission from Andor.

1. View each channel of the Maximum projections and hit “Command + M” or navigate ‘Analyze’ è ‘Measure.’
2. Repeat for each channel and assign these values to the following variables depending on the channel the value was taken from:

a. *X_maxZ OPC_*
b. *X_maxZ pSymDig-mCherry_*

*Whereby OPC is the channel with NG2 or the reporter line and pSynDig-mCherry is the mCherry fluorophore from the pSynDig construct*.

3. Save these values in an Excel file.
59. Based upon these variables, calculate the threshold values (T) that will be used to create surfaces for the cell and pSynDig markers in Imaris using the following formulas:

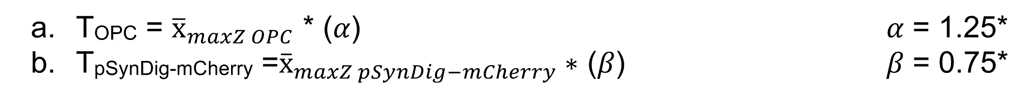

*NOTE: Both *α* and *β* are arbitrary values that can be adjusted by the investigator during the initial data analysis and should be based off the relative signal to noise of the experiment. Once set, these values should be kept consistent throughout the rest of the data analysis*.

***\*The values stated above are EXAMPLES and should not be used blindly.*** CRITICAL STEP At the beginning of the dataset, optimization of *α* and *β* is suggested. Optimize *α* by identifying a clearly defined NG2 cell stain and a less defined NG2 cell stain, and trying different *α* values to optimize the variable to be effective for both cases. Optimize *β* by doing the same but for the pSynDig-mCherry signal.

### Imaris pSynDig Analysis – OPC Surface Creation

60. Navigate back to Imaris and toggle off all the channels except for the channel representing the OPC (NG2).
61. Select the Surface icon (Figure 6b, teal box) to open the Creation Wizard (Figure 6b, teal arrow). When prompted, choose the OPC channel and select “Absolute Value” as the threshold (Figure 6bi)

a. Use the calculated threshold value for OPCs (TOPC) to set the lower threshold.
b. Click through the rest of the wizard, deleting any suggested filters. Finish the surface creation and rename the surface to be ‘All OPCs.’
c. OPTIONAL: Save Parameters for ‘All OPCs’ surface creation by navigating to the ‘Creation’ tab (wand) and select ‘Store Parameters for Batch’ and save.
62. Use the Orthogonal slicer (Figure 6bi, dashed teal box) and selection tool to choose the surfaces that represent a single cell of interest. Hold down command/control while selecting the disconnected OPC surfaces and use the arrow keys to move the orthogonal slicer through the Z-stack. Confirm that the elements reconstructed in the surface are representative of the OPC by comparing the reconstruction to the NG2 fluorescence channel. TROUBLESHOOTING

a. Select ‘Edit’ (pencil icon) and click on ‘Duplicate’ to duplicate the surface (Figure 6bii). Rename this surface to “OPC 1”.
b. Use the selection tool to select all parts of the OPC 1 surface and go to ‘Edit’ and click ‘Unify’ to unify the individual parts of the surface into one.
63. Repeat step 8 for all cells in the image within Layer 4 of V1. Each cell should have its own individual surface.
64. Repeat this entire process (Steps 60-63) for all images. For step 61, you may select your saved ‘All OPCs’ creation parameters under the ‘Favorite Creation Parameters’ dropdown CRITICAL STEP Make sure that you change the TOPC to the appropriately calculated threshold for each image.

### Imaris pSynDig Analysis – pSynDig Surface Creation

65. Select the Surface icon (Figure 6b, teal box) from the Imaris Surface Object Menu to open the Wizard for surface creation. When prompted, choose the **mCherry channel** to create the new surface (Figure 6bi).
CRITICAL STEP When setting the threshold of the surface, make sure ‘local background subtraction’ is selected.

a. Set ‘Diameter of largest Sphere which fits into the Object:’ to 1.00 µm.
b. Use the calculated threshold value for the marker (TpSynDig-mCherry) to set the lower threshold.
c. Click through the rest of the wizard, deleting any suggested filters. Finish the surface creation.
d. Rename surface to represent the marker of interest.
e. **OPTIONAL**: Save Parameters for ‘All mCherry’ surface creation by navigating to the ‘Creation’ tab (wand) and select ‘Store Parameters for Batch’ and save.
66. Apply a filter to the marker surfaces to retain only the marker surfaces that are completely within the cell surface of interest.

a. Navigate using the Object Menu Bar to ‘Edit’ (pencil)è ‘Image Properties’ è ‘Geometry (Figure 6aiii) and look under the ‘Coordinates [µm]’ section to see image dimension parameters (Figure 6aiv).

i. Dx,y = voxel size of X and Y (should be the same)
b. Calculate the filter threshold:

i. T_filter_ = -1 * (Dx,y * 2)
ii. *Note: The units of Tfilter should be some unit of length (nm or µm)*
c. With your pSynDig-mCherry surface selected, go to the Object Menu Bar (Figure 6biii) and select ‘Filter’ (funnel icon). Add a new filter by clicking on the ‘+’ icon. From the drop box menu, select the ‘Distance from nearest surface; surface = *name of OPC surface.*’
d. Use the calculated filter threshold (Tfilter) to set the **upper threshold** of the distance filter. Toggle off the lower threshold.
e. Rename surface to represent the marker of interest (i.e., pSynDig-mCherry within OPC 1)
67. Select all the pSynDig-mCherry surfaces created from this approach (within OPCs) and go to ‘Edit’ è ‘Unify’ (Figure 6bii). TROUBLESHOOTING
68. Repeat this entire process (Steps 1-2) for all images. For step 1, you may select your saved ‘ALL mCherry’ creation parameters under the ‘Favorite Creation Parameters’ dropdown menu. CRITICAL STEP Make sure that you change the TOPC and TpSynDig-mCherry to the appropriately calculated thresholds for that image.

### Imaris pSynDig OPC Quantification

69. Save progress and navigate to the Arena via the Arena and Batch Menu (Figure 6a, purple arrow)
70. Select the images that have been analyzed.
71. Select ‘Vantage’ from the Arena and Batch Menu (Figure 6c).
72. Under ‘Surpass Objects and Labels’ select the boxes representing the pSynDig-mCherry surfaces (if an image contains multiple cells, each cell should contain an individual pSynDig-mCherry surface that is uniquely named) (Figure 6ci).
73. Select ‘2D View’ from the Arena and Batch Menu (Figure 6c).
74. Under ‘Plot Type’ choose ‘Sum Intensity *mCherry*’ and ‘Sum Intensity *eGFP’* (Figure 6cii). CRITICAL STEP Make sure to note beforehand which channel represent mCherry and eGFP prior to this step to avoid confusion.
75. Navigate to ‘Plot Numbers Area’ è ‘Detailed.’ This will display the Sum Intensities of mCherry and eGFP within the pSynDig-mCherry surfaces. Select the save icon and save the data as an excel sheet. (Figure 6ciii). TROUBLESHOOTING
76. Open the spreadsheet of the Sum intensities data in Excel.
77. Normalize the eGFP and the mCherry signals of an individual cell to the average mCherry signal of the dataset.
78. Copy and paste these normalized eGFP and mCherry intensities into GraphPad Prism (or a similar program) for statistical analysis.
79. Conduct paired parametric tests for Gaussian datasets and nonparametric tests for datasets in which a parametric analysis is inappropriate. Before unblinding and finalizing analyses, remove outliers using the ‘Identify Outliers’ function in GraphPad (using the robust regression and outlier removal (ROUT) method with Q (maximum desired false discovery rate) = 1%).

a. Conduct a paired test to compare the eGFP signal within an OPC to its respective mCherry signal from that same cell.

CRITICAL STEP Choosing the correct statistical test is non-trivial and consultation with a biostatistician or with online guides (i.e., GraphPad guides) is vital to conducting appropriate statistical tests on biological datasets.

### Imaris OPC Engulfment Quantification

Note: An engulfment score can be obtained through the mCherry surfaces as they are representative of the VGLUT2+ inputs from the dLGN (Figure 6biv). Validation of pSynDig expression and proper dLGN transduction through co-staining with VGLUT2 is recommended prior to quantification.

80. Follow steps 69-72 as stated above.
81. Select ‘1D View’ From the Arena and Batch Menu.
82. Under ‘Plot Type’ choose ‘Volume.’ And navigate to ‘Plot Numbers Area’ è ‘Detailed’ to see the Volumes of the pSynDig-mCherry surfaces. There should be one volume for all the pSynDig-mCherry surfaces in a single OPC. Select the save icon and save the data as an excel sheet. TROUBLESHOOTING OPTIONAL: This analysis can also be done without pSynDig but with just staining tissue sections for NG2 and VGLUT2 (Table 1).
83. Repeat steps 81-82 but instead select the boxes under “Surpass Objects and Labels’ representing the OPC surfaces to obtain the individual OPC volumes.
84. Normalize each pSynDig-mCherry volume to its respective OPC volume to obtain an engulfment score for that individual OPC. Copy these scores into GraphPad Prism.
85. When comparing the engulfment scores for two different conditions, follow the logic described in step 79 but conduct a non-paired statistical test.

### Timing SECTION A

Steps 1–14, General Pre-surgery preparation: 30 minutes. Steps 15–25, Stereotaxic Injection: 1-2 hours

Steps 26–27, Post-surgery: 2-3 weeks

Steps 28–33, Animal sacrifice and perfusion for immunofluorescence staining: 2-3 days. Steps 34-39, Cryosectioning: 3-5 hours.

Steps 40-51, Immunofluorescence staining: 2 days.

Steps 52-53, Confocal Image Acquisition: 3 to 9 hours.

Steps 54-85, Imaris Analysis: 6 hours - 4 days.

## SECTION B – Flow cytometry protocol

### Materials

#### Animals

Only mice with C57Bl/6 backgrounds have been validated for this protocol, but other stains most likely can be used as well. ▲ CAUTION Any experiments using animals must comply with National Institutes of Health (NIH) guidelines on animal care. All protocols were approved by the Institutional Animal Care and Use Committee (IACUC) at CSHL.

### Reagents

#### Flow cytometry reagents

CRITICAL The reagents and antibody panel (Table 1) used here are specific for OPC staining.

- ArcReactive Amine beads (LIVE/DEAD) (Thermo Fisher, cat. no. A10628)
- UltraComp beads (antibodies) (Thermo Fisher, cat. no. 01-2222-41) CRITICAL In this protocol, we are using Compensation beads for the single stained control due to the limited availability of cellular material. However, the beads can be replaced by cells for generating the single stained control in case the cell population is abundant.
- 1X HBSS / 10X HBSS (Thermo Fisher, cat. no. 14175095 / 14185052)
- PBS 1X, pH 7.4 (Thermo Fisher, cat. no. 10010049)
- Percoll (Sigma-Aldrich, cat. no. GE17-0891-02)
- LIVE/DEAD aqua staining for viability (Thermo Fisher, cat. no. L34957)
- D-(+)-Glucose powder (Sigma-Aldrich, cat. no. G8270)
- Molecular Biology Grade Water, DEPC Treated, 1 L (Ricca, cat. no. R9145000)
- BSA AlbuMAX II Lipid Rich BSA powder (Thermo Fisher, cat. no. 11021029)
- EDTA 0.5M pH8.0 (Sigma-Aldrich, cat. no. 03690-100ML)
- HEPES 1M (Sigma-Aldrich, cat. no. 15630080)
- Accumax (Thermo Fisher, cat. no. SCR006)
- eBioscience™ Foxp3 / Transcription Factor Staining Buffer Set (Thermo Fisher, cat. no. 00-5523-00)
- 16% (vol/vol) Paraformaldehyde solution (electron microscopy grade; Thermo Fisher/Electron Microscopy Sciences, cat. no. 15710)

#### Equipment General equipment

- Watson Marlow 205CA4 Channel pump with Pump Pro MPL (Boston Laboratory

Equipment cat. no. BLE2000180) CRITICAL This system can be replaced with other perfusion pumps

- PVC Tubing (1/16 x 1/8 in) (Sigma-Aldrich cat. no. Z280348)
- 22-gauge needle with tip cut off (VWR cat. no. BD305156)
- Mini Dissecting Scissors, 8.5 c (World Precision Instruments (WPI) cat. no. 503667)
- Operating Scissors straight 11.5 cm (WPI cat. no. 501753)
- Dumont Tweezers #5 (WPI cat. no. 501985)
- Dressing Forceps (WPI cat. no. 500363)
- Round end spatula
- In-house vacuum line or vacuum pump
- Liquid aspirator setup
- Polypropylene FACS tubes 12 x 75 mm – 5 mL (VWR, cat. no. 60818-576)
- Falcon® tubes – 15 mL (VWR, cat. no. 21008-918)
- Falcon® tubes – 50 mL (VWR, cat. no. 21008-951)
- Sterile Falcon® Cell Strainers 70 µm (VWR, cat. no. 21008-952)
- Corning® tube/bottle top vacuum filtration system (Sigma Aldrich, cat. no. CLS430320-12EA/CLS431205-12EA)
- Benchmark Scientific Roto-Mini Plus Tube Rotator (Stellar Scientific, cat. no. BS-RTMNI-2)
- Swinging-bucket centrifuge (Beckman coulter, model no. B99517 or equivalent)
- Mini Vortex mixer (VWR, cat. no. 10153-838)
- Flow cytometer (BD Biosciences, model no. BD LSR DualFortessa) or similar

#### Data analysis software

- Analysis tools: CytoExploreR^23^ (v. 1.1.0, Dillon Hammill) (https://dillonhammill.github.io/CytoExploreR/index.html) or FlowJo™ Software (v. 10.8.1 for Mac, BD Life Sciences) (https://www.flowjo.com/solutions/flowjo) for flow cytometry data analysis.
- Flow cytometry panel design tool such as FluoroFinder (https://fluorofinder.com).
- BD FACSDiva (v. 6.0, BD Biosciences)

### Reagent setup

#### FACS tubes with 0.5X Accumax enzyme

Prefill the propylene FACS tubes with 1:1 of HB and 1X Accumax (500 µL of each) before starting the experiment. If working with 4 samples, prepare 4 FACS tubes filled with 0.5X Accumax. Store the tubes in the 4°C until use.

#### PBS pH 7.4 or Saline 0.9%

Place the PBS pH 7.4 or Saline 0.9% in the fridge a day before the experiment. If using saline, mix 9 g of NaCl in 1 L of water. Prepare fresh and keep the reagents ice-cold.

#### Homogenization buffer (HB)

Mix 1.5 mL of 1M HEPES (150 mM final concentration), 10 mL of 10X HBSS (1X final concentration), 5 g of Glucose (5 % final concentration), 1 g of BSA (1% final concentration), 400 µL of 0.5 M EDTA pH 8.0 (2 mM final concentration) and fill up to 100 mL with DEPC water and filter using 0.22 µm top vacuum filtration system. Prepare fresh and keep the homogenization buffer ice-cold.

## 40% Isotonic percoll

Prepare 20 mL of isotonic percoll by mixing 18 mL of percoll with 2 mL of 10X HBSS. Next, mix 15.5 mL of isotonic percoll with 19.5 mL of room temperature (RT) 1X HBSS. Prepare fresh and keep the Isotonic percoll solution at RT until use.

### Blocking solution

Prepare blocking solution at a 1:50 dilution. If starting with 4 whole cortices (4 mice), 250 µL of blocking solution will be needed. Prepare an eppendorf tube with 250 µL 1X HBSS + 5 µL of anti-CD16/32. Prepare fresh right before use.

Extracellular Antibody and Viability dye solution (Pre-fixation/permeabilization) Prepare the extracellular antibody solution at a 1:50 dilution and the viability dye at a 1:500 dilution. 250 µL of extracellular antibody solution is needed for 4 whole cortices worth of cells. Prepare a tube with 250 µL 1X HBSS + 5 µL of each antibody + 0.5 µL of viability dye according with the list below:

a. 5 µL of anti-CD140a-Pe-Cy7
b. 5 µL of anti-A2B5-AF488
c. 0.5 µL of LIVE/DEAD Aqua.

Prepare fresh, right before use. CRITICAL The same volume of blocking and extracellular antibody/viability dye solution will be combined after the blocking step, making the final antibody concentration 1:100 for the extracellular antibodies and 1:1000 for the viability dye.

### Intracellular Antibody solution (Post-fixation/permeabilization)

Prepare the intracellular antibody solution at a 1:100 dilution. 500 µL of intracellular antibody solution is needed for 4 whole cortices worth of cells. Prepare an eppendorf tube with 500 µL HB + 5 µL of each antibody according to the list below:

a. 5 µL of anti-SYNAPSIN1-AF647

Prepare the solution fresh, right before to use.

## 1% (vol/vol) PFA

Prepare the solution according to the number of samples you are running (0.5 mL/ sample). Mix 1 part 16% PFA with 15 parts 1X HBSS. For 4 samples, use 156 µL of 16% PFA and 2344 µL of 1XHBSS. ▲CAUTION Toxic reagent. Always handle with gloves and avoid eye and skin contact. Handle reagent in hood with high airflow. Only use with designated PFA tools.

### Fix/Perm staining buffer (eBioscience™ Foxp3 / Transcription Factor Staining Buffer Set)

Prepare fresh Fix/Perm buffer according to the manufacturer’s instructions. Briefly, mix 1 part of Fix/Perm buffer with 3 parts of Permeabilization Diluent right before use. For 4 whole cortices worth of cells, use 625 µL of Fix/Perm buffer and 1875 µL of Permeabilization Diluent.

### Equipment setup

### Swinging-bucket centrifuge

Cool down the centrifuge to 4°C before use.

### Flow cytometer

Set up the machine according to the flow cytometry facility or manufacturer’s instructions.

#### Procedure

### Preparation of reagents and perfusions ● 30 min

1 Set up bench and materials for dissection and collection (this can be prepared the day before). Wash the tools and keep the materials ice-cold until use. Prepare 2 ice buckets: one for dissection and collection of the material and another for the staining protocol.
2 Prepare the FACS tubes prefilled with 0.5X Accumax (see <u>REAGENT SETUP</u>).

### Tissue collection ● 1h

3 Anesthetize mice with a method suitable for perfusion. We recommend anesthetization with isoflurane followed by transcardial perfusion with ice-cold 1X PBS while maintaining anesthesia. Collect the brains and dissect the cortices in PBS on ice.

### Enzyme digestion ● Overnight (ON)

4 Transfer the cortices to a FACS tube prefilled with 1 mL of 0.5X Accumax enzyme. Collect one whole cortex/tube or, if working with pooled material, use a maximum of 4 whole cortices/tube and increase the volume of 0.5X Accumax to

## 1.5 mL

5 Using a round end spatula, slowly chop the cortices to obtain a suspension of cortical pieces of about 2-3 mm in size.
6 Wrap the lid with parafilm and incubate overnight at 4°C in a rotating mixer.

### Homogenization ● 45 min

CRITICAL STEP For each sample, prepare a set of P1000 tips and P200 tips cut about 1 and 0.5 cm short, respectively.

CRITICAL STEP The homogenization step should not take longer than 5 min per tube (about 1 min per tip).

7 After overnight incubation, with the tissue still in the FACS tube, homogenize the sample using a P1000 pipette with 1 cm short-cut tip. Gently pipette the homogenate up and down until the suspension moves freely in the tip. TROUBLESHOOTING
8 Repeat step 7 using a P1000 pipette with 0.5 cm short-cut tip.
9 Repeat step 7 using a P1000 pipette with an uncut tip.
10 When all tissue is homogenized using a P1000 uncut tip, spin down tubes for just a quick spin at 300 g for 20 sec, 4°C to bring remaining chunks to the bottom.
11 Transfer the supernatants of each to a clean 15 mL conical tube, leaving about 400 µL in the original tube.
12 Homogenize the remaining tissue fragments in the original tubes using a P200 similarly to steps 7-9.
13 Transfer the remaining supernatant to the 15 mL conical tube. Wash the original tubes with 2 mL of HB and combine with the suspension in the 15 mL conical tubes.
14 Spin down the samples at 300 g for 5 min, 4°C and gently discard the supernatant using the vacuum aspirator.
15 During the centrifugation, prepare the 40% isotonic percoll solution. (See <u>REAGENT SETUP</u>). About 50 mL is needed for a total of 4 samples (8 mL/cortex).
16 Discard the supernatant and proceed to the percoll separation step.

### Percoll separation ● 30 min

CRITICAL STEP The isotonic percoll and 1X HBSS solutions should be at RT. CRITICAL STEP Set the centrifuge to room temperature at this step. After centrifugation, set it back to 4°C for the following steps.

17 Gently resuspend each sample pellet in 1 mL of 40% Isotonic percoll using a P1000 pipette.
18 Add another 7 mL of 40% Isotonic percoll to each sample and slowly mix by inversion.
19 Centrifuge at 600 g for 25-30 min, 20°C. Make sure to set up the centrifuge for minimal brake (Acc. 5 and Decel. 1). CRITICAL STEP After centrifugation, it is expected to observe cellular debris and myelin in the upper layer of the supernatant, and OPCs, along with other cells of similar density, clustered at the bottom of the tube. TROUBLESHOOTING
50 Using a vacuum aspirator, carefully remove the myelin and supernatant leaving about 300 µL in the tube.
21 Gently resuspend the pellet in 1 mL of HB with a P1000 and transfer to a new 15 mL conical tube.
22 Fill the tubes with about 5 mL HB and spin down at 300 g for 5 min, 4°C, max Acceleration and max Deceleration.
23 Gently remove the supernatant and resuspend the pellet in 1 mL of cold 1X HBSS to remove proteins. Split one of the sample tubes into 2 new 15 mL conical tubes to obtain the Unstained and FMO-SYN1 controls, respectively. Spin as in step 22.

### Cell surface staining ● 45 min

CRITICAL STEP In order to perform LIVE/DEAD staining, cell pellets should be washed in 1X HBSS to avoid any interference from proteins.

CRITICAL STEP Most of the fluorophores are susceptible to photo bleaching resulting in a loss of fluorescence signal. Avoid over-exposure of the stained samples to light sources.

24 Resuspend each tube with 50 μl of the blocking solution (see REAGENTS SETUP) and incubate for 10 min on ice.
25 After the incubation step with the blocking solution, add 50 μl of the antibody solution (see REAGENTS SETUP) to each tube, with exception of the unstained control which should have 50 μl HB added. In this step you should have a final volume of 100 µL/tube. Incubate 20-30 min on ice protected from the light.
26 During the antibody incubation, prepare the beads for compensation according to the manufacturer’s instructions.
27 Wash the tubes with 5 mL of cold 1XHBSS and spin as in step 22. During the centrifugation, prepare 1% PFA solution (See REAGENT SETUP) and keep it at RT.

### Fixation and Permeabilization ● 50 min

CRITICAL STEP Setup the centrifuge to spin at 1000 *g* before the fixation and permeabilization step. Running the samples at a lower speed might cause cell loss.

28 Resuspend the pellets in 500 µL of 1% PFA and incubate at RT in the dark for 10 min.
29 Wash the cells by adding 5 mL of 1X HBSS buffer per tube. Spin down the samples at 1000 *g* for 5 min at RT. During the centrifugation, prepare the fix/perm buffer (See REAGENT SETUP). TROUBLESHOOTING
30 Resuspend the pellet in 500 µL of diluted fix/perm buffer and incubate at RT in the dark for 30 min.
31 Wash the cells by adding 5 mL of HB per tube. Spin down the cells at 1000*g* for 5 min, 4°C.

### Intracellular staining ● 40 min

32 Resuspend the pellet of each sample in 100 µL of intracellular antibody (See REAGENT SETUP), except for the control, unstained, and FMO-SYN1 samples, to which 100 µL of HB should be added before incubating all samples for 30 min on ice protected from light.
33 Wash the cells by adding 3 mL of HB per tube and spin down the cells at 1000 *g* for 5 min, 4°C. PAUSE POINT Either proceed to flow cytometry acquisition or optionally store at 4°C in the dark for a maximum an overnight. CRITICAL STEP If using tandem dye conjugated antibodies proceed immediately.

### Flow cytometry data acquisition ● 1-3 h

34 Resuspend each pellet in 1 mL of HB and filter through a 35 μm cell strainer in a polypropylene FACS tube. Make two additional washings with 1 mL of HB each to collect as many cells as possible when transferring to the FACS tubes.
35 Spin down the cells at 1000 *g* for 5 min at 4°C. Remove supernatant leaving about 150 µL, resuspend the pellet in 350 µL HB and bring the samples to the flow cytometry facility for the data acquisition.
36 Run the experimental and control tubes in a flow cytometer and acquire the .FCS files using flow cytometry software for further analyses. CRITICAL STEP Manual and automated compensation can be generated and linked to the .FCS files during acquisition. Alternatively, the compensation .FCS files can be acquired separately, and the samples can be compensated after acquisition. CRITICAL STEP ArcReactive Amine and UltraComp beads have different sizes. Ensure to adjust the side and forward scatter lasers in order to have all the beads and cells in the same plot. If not possible, prioritize the alignment of the cells. TROUBLESHOOTING

### Data analysis ● 1 h

37 Import all .FCS files from step 36 into the flow cytometry analysis software of choice and transform the cytometry data using *Logicle* or *Bioex* transformation. CRITICAL STEP The transformation of flow cytometry data is essential for proper visualization of the cytometry data. The most commonly used transformations are Logicle and Bioex transformations^24^, as they offer good visualization of both discrete negative and positive values. It should be noted that each parameter is dependent on the data obtained at the time, and the transformations should be optimized by the user for better visualization.
38 Start by gating the cells of interest using side and forward scatter lasers (SSC-A versus FSC-A). Only exclude those events that you are sure to not be of interest (Figure 5a).
39 Subsequently, gate on Singlets 1 (diagonal of FSC-H versus FSC-A) and Singlets 2 (SSC-W versus SSC-A) (Figure 5a).
40 Next, gate the negative events by using the LIVE/DEAD Aqua laser for the exclusion of dead cells (Figure 5a). CRITICAL STEP The use of viability dye is essential for a clear interpretation of the results since dead cells can bind non-specifically to the antibody.
41 Identify the OPC population by gating the positive events for both A2B5 AF488 and CD140a PE-Cy7 lasers.
42 At this point, it is useful to apply backgating to analyze the effectiveness of the gating strategy (Figure 5b). This technique will reveal whether all cells of interest, OPCs in this case, have been correctly grouped for downstream analysis.
43 Once the OPCs are isolated from the rest of the cellular events, analyze the target protein using the A2B5 AF488 versus SYNAPSIN1 AF647 lasers.
44 Create an overlay plot of the FMO-SYN1 control over the experimental samples to define correctly the positive events for SYNAPSIN AF647 laser to be gated and the different populations. CRITICAL STEP The FMO-SYN1 control was used here to identify all the positive events for SYN1. To discriminate for other populations, one can explore contour or densities plots to better depict a population.
45 Generate the figures and export all the data (counts, mean fluorescence intensity, population percentage, etc.) regarding the experimental and control samples as .CSV files for future statistical analyses. TROUBLESHOOTING

#### Timing SECTION B

Steps 1–3, Preparation of the reagents, perfusion, and tissue collection: 1 h 30 min. Steps 4–6, Enzyme digestion: 16 h or ON.

Steps 7–16, Homogenization: 45 min.

Steps 17–23, OPCs isolation: 40 min.

Steps 24–33, Cell surface staining, Fixation/permeabilization and Intracellular staining: 2 h 15 min.

Steps 34–36, Flow cytometry data acquisition: 1-3h. Steps 37–45, Data analysis: 1h.

### Anticipated results

Recent studies have revealed OPCs to be highly dynamic cells with crucial functions in development, homeostasis, and disease^25^. One important role that OPCs play in the brain is to eliminate synapses through phagocytic engulfment. Here, we detail two methods for quantifying synapse engulfment by OPCs highlighted in a previous publication^9^. The first method employs a viral fluorescent sensor of synaptic digestion (pSynDig) yielding two quantitative outputs reflecting the amount of synaptic material engulfed by OPCs. The first output is a ratiometric measurement of the sum intensities of the mCherry and eGFP signals found within OPCs. pSynDig-expressing inputs that are in the process of degradation lack eGFP fluorescence while inputs that remain intact are labeled with both mCherry and eGFP. Thus, if a given OPC is in the process of degrading synaptic inputs labeled with the pSynDig construct, the normalized eGFP signal within that cell is anticipated to be significantly lower than the respective mCherry signal. As reported previously, co-staining the tissue for markers of mature phagosomes, such as LAMP2, can be employed to validate that inputs lacking eGFP fluorescence reside within acidic intracellular compartments^9^. Overall, if OPCs are engaged in engulfing synaptic material, the synapses inside OPCs are expected to have a lower eGFP:mCherry ratio than synapses outside OPCs.

The second measurement derived from the pSynDig analysis is an engulfment score which is obtained through the quantification of synaptic material within OPCs as analyzed in Imaris. The engulfment score represents the volume of engulfed synaptic material present within an OPC, as measured by a distance filtering-based method, normalized to the volume of the reconstructed OPC. OPC engulfment scores across different conditions can be compared to determine differences in the engulfment capacity of OPCs across numerous contexts.

The second method described here is a simple and efficient flow cytometry protocol for high-throughput analysis of synaptic engulfment by OPCs. In this approach, the detection of positive events for SYN1 is expected, indicating the presence of synaptic material within OPCs. A majority of OPCs are expected to contain moderate levels of SYN1, while a smaller subset of OPCs exhibits high levels of SYN1 and a third set of OPCs do not contain SYN1 beyond FMO control levels (Figure 5c and 5d).

Metadata information (population frequencies, counts, MFI, etc.) should be used to understand the frequency of these populations, infer the amount of SYN1 inside of the cells and to identify possible alterations in engulfment occurring across different stages of development or under pathological conditions. It is also important to mention that non-cortical brain regions may have different cellular dynamics, requiring additional validations prior to data interpretation.

### Adaptations and improvements

We predict that the approaches described here will continue to undergo evolution and refinement as more investigators adopt the strategies. To this end, we have identified some potential aspects of the protocols that may be optimized or improved over time. For example, in the imaging-based assay, the NG2 antibody used to label OPCs works best under conditions of light fixation which are often not optimal for other antibodies^26,27^. In the future, better results could be achieved through the use of antibodies against a second OPC marker, PDGFRα, or by using a transgenic line that labels OPCs with a cell-filling fluorophore. We have had good success with a goat anti-PDGFRA antibody (R&D Systems, cat. no. AF1062) and a mouse line derived from a cross between B6.Cg-Tg(Cspg4-cre/Esr1*)BAkik/J (Jackson Laboratory strain code 008538) and the tdTomato fluorescent reporter line ROSA-CAG-LsL-tdTomato (Jackson Laboratory strain code 007914). A second adaptation which could be possible in the future is the use of the pSynDig construct, and similar constructs with improved pH-sensitivity, to analyze synapse engulfment by OPCs *in vivo*. This approach would require the development of OPC markers that fall outside of the red/green range. Also, in Imaris, OPC cell reconstruction could be improved by using a Fiji plugin called Labkit, a machine-learning pixel classifier. Experienced investigators can train this function to identify and reconstruct the entirety of the OPC (including un-stained intracellular compartments and thinner processes) as a whole. Consultation with the Imaris support desk is suggested for implementing these improvements.

For the flow cytometry-based approach, it is worth noting that we have limited this manuscript to the analysis of synaptic material by detecting the presynaptic protein SYN1, but this protocol can be adapted for the analysis of different synaptic proteins such as VGLUT1, VGLUT2, and SNAP25^9^. Furthermore, the methodology used here for the isolation of OPCs allows for the joint analysis of different cell types. To this point, it is useful to include microglial cells in the analysis of the data, as it is possible to evaluate the phagocytic efficiency of OPCs when directly compared to cells that also play a fundamental role in synaptic refinement and modulation of neuronal connections through phagocytosis. Overall, we expect the approaches described here to provide significant new insights into the roles of OPCs in the developing and mature brain.

## Author contribution statement

J.A.K., A.M.X., and L.C. wrote the paper. For the material, reagents, and protocol sections, the components describing the imaging-based approach were written by J.A.K. and the components related to flow cytometry were written by A.M.X. J.A.K., A.M.X., and L.C. contributed to the introduction and discussion sections. A.F. designed and produced the pSynDig construct, and A.F., J.A.K., and Y.A. contributed to optimizing the pSynDig engulfment assay analysis. All figures were created by J.A.K., A.M.X., and L.C.

## Acknowledgements

We would like to acknowledge and thank the following individuals for their contributions: Uma Vrudhula for contributions to early imaging-based engulfment protocols, Chris Kang for providing an image dataset of OPCs, Anne-Sarah Nichitiu for previously validating the pSynDig construct, Dr. Pamela Moody from the CSHL Flow Cytometry Core, Dr. Erika Wee from the CSHL Microscopy Core, and Dr. Matthew J. Gastinger from Imaris/Andor. This work was supported by the following funding sources (to L.C.): R00MH120051, DP2MH132943, R01NS131486, Rita Allen Scholar Award, McKnight Scholar Award, Klingenstein-Simons Fellowship Award in Neuroscience, and a Brain and Behavior Foundation NARSAD grant.

